# Cnpy1 is a candidate endoplasmic reticulum chaperone of Vomeronasal type 2 GPCRs

**DOI:** 10.1101/2025.10.06.680635

**Authors:** G V S Devakinandan, Abdul Rishad, Nandana Nanda, Syed Dastagir Hussain, Sishir Subedi, Adish Dani

## Abstract

Mouse vomeronasal sensory neurons are continuously generated from stem cells and differentiate to express either V1R or V2R G-protein coupled receptors (GPCRs), along with their respective Gαi2 or Gαo G-protein subunits. We previously reported that Gαo-type neurons exhibit elevated expression of endoplasmic reticulum (ER) chaperones and a distinctive hypertrophic, gyroid ER architecture, suggesting specialized proteostatic demands. Here we identify a transcript for the mouse *Cnpy1* gene that yields full-length Cnpy1 protein selectively expressed in and localized to the ER of Gαo neurons. Immunoprecipitation coupled with mass spectrometry revealed that Cnpy1 associates specifically with V2R GPCRs and multiple ER chaperones. *Cnpy1* deletion resulted in mice that were deficient in Gαo neuronal activation upon exposure to vomeronasal stimuli and a marked reduction in male-male aggressive behavior. In the absence of Cnpy1, Gαo neurons develop normally till birth but undergo selective, progressive apoptosis during postnatal development. Unexpectedly, Cnpy1-null vomeronasal neurons displayed neither an obvious unfolded protein response nor defects in V2R GPCR traffic to dendritic tips, indicating that Cnpy1 is required for V2R assembly or functional maturation but dispensable for their ER export. Together, these findings identify Cnpy1 as a previously unrecognized component of an ER chaperone complex that is essential for Gαo neuron signaling and survival.

## Introduction

Animals depend on a highly diversified chemosensory system to sense their environment, seek food as well as social and inter-species interactions. The vomeronasal or accessory olfactory system, comprised of a peripheral vomeronasal organ (VNO) specializes in detection of pheromones, kairomones and such chemo-signals that elicit innate stereotyped behaviors. Several vertebrates, including rodents, have evolved an elaborate VNO molecular organization, with major classes of G-protein coupled receptor (GPCR) families - vomeronasal type-1 (V1R), type-2 (V2R), or formyl-peptide receptors (FPR) (1) being expressed in spatially segregated sensory neurons within the neuroepithelium, along with the G-protein subunits, Gαi2 or Gαo. VSNs arising from a regenerating stem cell population, differentiate and migrate within the neuroepithelium to take on the identities of Gαi2 expressing V1R or Gαo expressing V2R neuronal subtypes. During this process, VSNs transition through distinct stages of progenitor, immature and mature cells, with choreography of cell autonomous and non-cell autonomous differentiation factors. Thus, in addition to serving as an important model system to understand the neurobiology of innate behaviors, the VNO is also a model to study neuronal development and diversification, a process that occurs throughout the lifespan of the organism.

While few ligands that activate V1R or V2R expressing neurons have been identified by calcium or immediate early gene activation studies (2–5) the downstream signaling and cell biology of Gαi2 and Gαo neurons remain poorly understood. One of the hurdles has been the inability to reconstitute functional expression and signaling of VRs in heterologous cells, due to GPCR misfolding and retention inside the endoplasmic reticulum (ER), a feature shared with olfactory receptors (OR). Given that OR, V1R, V2R GPCRs are evolutionarily distinct, their expression, maturation, signaling and cell biology may necessitate neurons expressing these GPCRs to diversify in their cellular and molecular components, while sharing certain features. Indeed, earlier studies have identified genes such as *H2-Mv* that are selectively co-expressed with V2Rs and may play a role in receptor function (6, 7). Several examples exist of specific factors, including ER chaperones that are required for the maturation and targeting of sensory GPCRs. The protein ODR-4, is required for ciliary localization of ORs in C.elegans (8), chaperones NinaA, Calnexin and XPORT are essential for Drosophila Rhodopsin GPCR maturation (9–11), receptor transporting proteins, RTP1 and RTP2 aid expression of mammalian ORs (12) and a VNO specific calreticulin variant, Calr4, has been shown to promote the maturation of V2Rs upon their expression in heterologous cells (13). Chaperoning of ORs and VRs is important for their proper targeting, but also for the survival of the sensory neuron (14).

In our earlier study (15), performing comparative transcriptomics of V1R expressing Gαi2 and V2R expressing Gαo neurons, we observed that mature Gαo neurons differ by significantly upregulating the expression of genes and proteins whose functions are associated with the endoplasmic reticulum (ER). Furthermore, Gαo neurons adopt a highly hypertrophic ER membrane organization, characterized by a cubic or gyroid membrane ultrastructure. These results suggest a fundamental cell biological divergence of Gαo neurons and prompted us to investigate the role of certain Gαo neuron specific upregulated genes. Here, we identify Cnpy1 as a VNO neuron specific protein, that is exclusively expressed in Gαo-VSNs and localized to the ER. The molecular and functional characterization of Cnpy1, including its absence, highlights the unexpected role of specialized ER proteins in Gαo neurons that maybe important for the function of V2R GPCRs, neuronal signaling and survival.

## Results

### Expression of *Cnpy1* in the VNO

In our recent study, we performed single cell transcriptomics of the mouse VNO and compared transcriptomes of Gαi2 with Gαo expressing neurons (15). This analysis resulted in the identification of several genes that are selectively upregulated in Gαo neurons. Amongst the Gαo enriched genes, we investigated Canopy1 (*Cnpy1*) in depth to understand its function. From our single cell RNA sequencing data, *Cnpy1* expression is restricted to immature and mature Gαo neurons, with no expression in Gαi2 neurons (**Figure 1A, 1B**).

**Figure 1.**
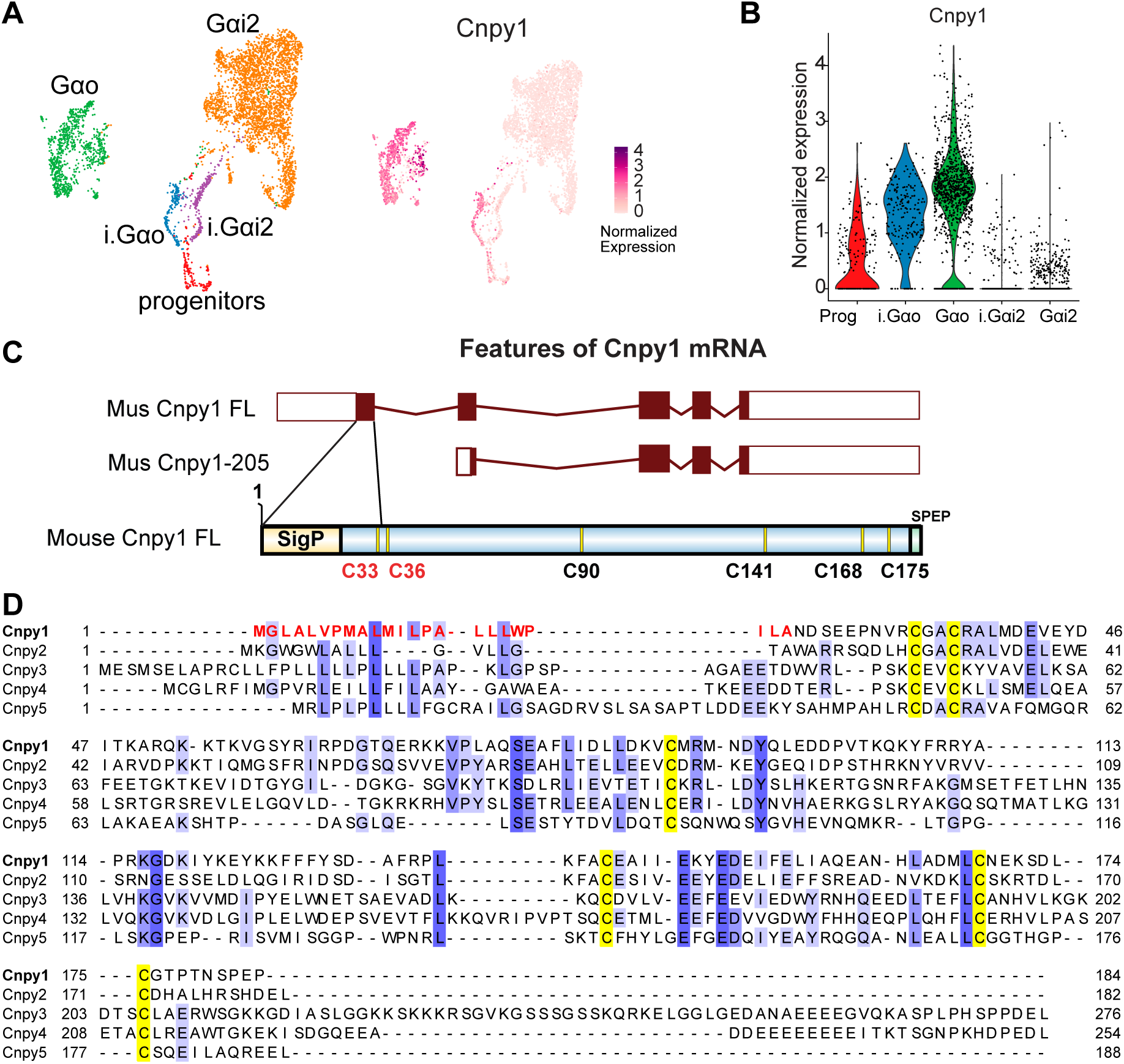
Description of the complete mouse Cnpy1 gene. **A**) UMAP (left, from Devakinandan et. al. 2024) and feature plot (right) showing Cnpy1 gene expression amongst vomeronasal sensory neuron clusters coded in different colors corresponding to progenitors (prog), immature Gαo (i.Gαo), immature Gαi2 (i.Gαi2), mature Gαo (Gαo), and mature Gαi2 neurons (G αi2). Cnpy1 expression is restricted to immature and mature Gαo neurons. **B**) Violin plot shows the distribution and extent of Cnpy1 expression in neuronal clusters. **C**) Schematic showing the transcript structure of full length Cnpy1 (Cnpy1-FL) and its comparison with the transcript from Ensembl database (Cnpy1-205, ENSMUST00000141601). Exons are represented as boxes, with coding regions shaded and untranslated regions as hollow boxes. The first exon in Cnpy1-FL contributes an upstream transla-tion initiation site for a longer ORF with a signal peptide (SigP, **Figure S1**) and two out of the six signa-ture cysteine residues comprising saposin-like domain. **D**) Alignment of Cnpy1 (Cnpy1-FL) with Cnpy2-5 mouse protein sequences. Conserved 6-cysteine signature is highlighted in yellow and puta-tive N-terminal signal sequence is highlighted in red.

*Cnpy1*, first described in zebrafish (16), belongs to the saposin-like family of proteins. Mammalian CNPY family members, Cnpy2-Cnpy5, homologs of Cnpy1, contain a characteristic six-cysteine saposin like protein (SAPLIP) domain (17, 18). However, mouse Cnpy1 in Ensembl (ENSMUSG00000044681) or NCBI (Gene ID: 26963) databases contains only 4 cysteines, lacking the complete SAPLIP domain and is considered to be a truncated protein with no function attributed yet. In order to confirm if mouse VNO expresses the truncated version of *Cnpy1*, we used VNO bulk RNAseq data (19) and assembled the transcriptome. Interestingly, we found that VNO expresses a *Cnpy1* transcript that contains an additional upstream exon. Translation of the assembled open reading frame indicates that the upstream translation start site from this additional exon contributes a putative signal sequence (1-23 amino acids) predicted by SignalP 6.0 (20) (**Figure S1**), as well as 2 cysteines that complete the SAPLIP domain (**Figure 1C**). Multiple sequence alignment of full length Cnpy1 protein sequence (Cnpy1-FL) with other mouse Cnpy proteins shows the conservation of six-cysteine signature (**Figure 1D**), indicating Cnpy1 may form a functional protein in the VNO.

### Cnpy1 is a Gαo neuron specific ER protein

We reasoned that additional exon of Cnpy1 consisting of an upstream translation start site and a signal sequence was not detected earlier because it may be only expressed in VNO and often VNO genes are poorly annotated in gene expression databases. To confirm if full length *Cnpy1* expression is specific to VNO we made cDNA from several adult mouse tissues and used these as templates for PCR-based detection. We amplified a region of the exon1-exon2 and exon2-exon3 of Cnpy1-FL. The tissue panel RT-PCR revealed that expression of *Cnpy1* mRNA is restricted mainly to VNO with a faint band observed in the main olfactory epithelium (MOE) (**Figure 2A**). To localize *Cnpy1* expression to olfactory regions, we performed RNA *in-situ* hybridization (RNA-ISH) with a Cnpy1 anti-sense riboprobe on sections of the VNO, MOE and accessory olfactory bulb (AOB) (**Figure 2B-D**). This experiment confirmed that *Cnpy1* mRNA is restricted to sensory neurons of the VNO and MOE, with no expression in AOB.

**Figure 2.**
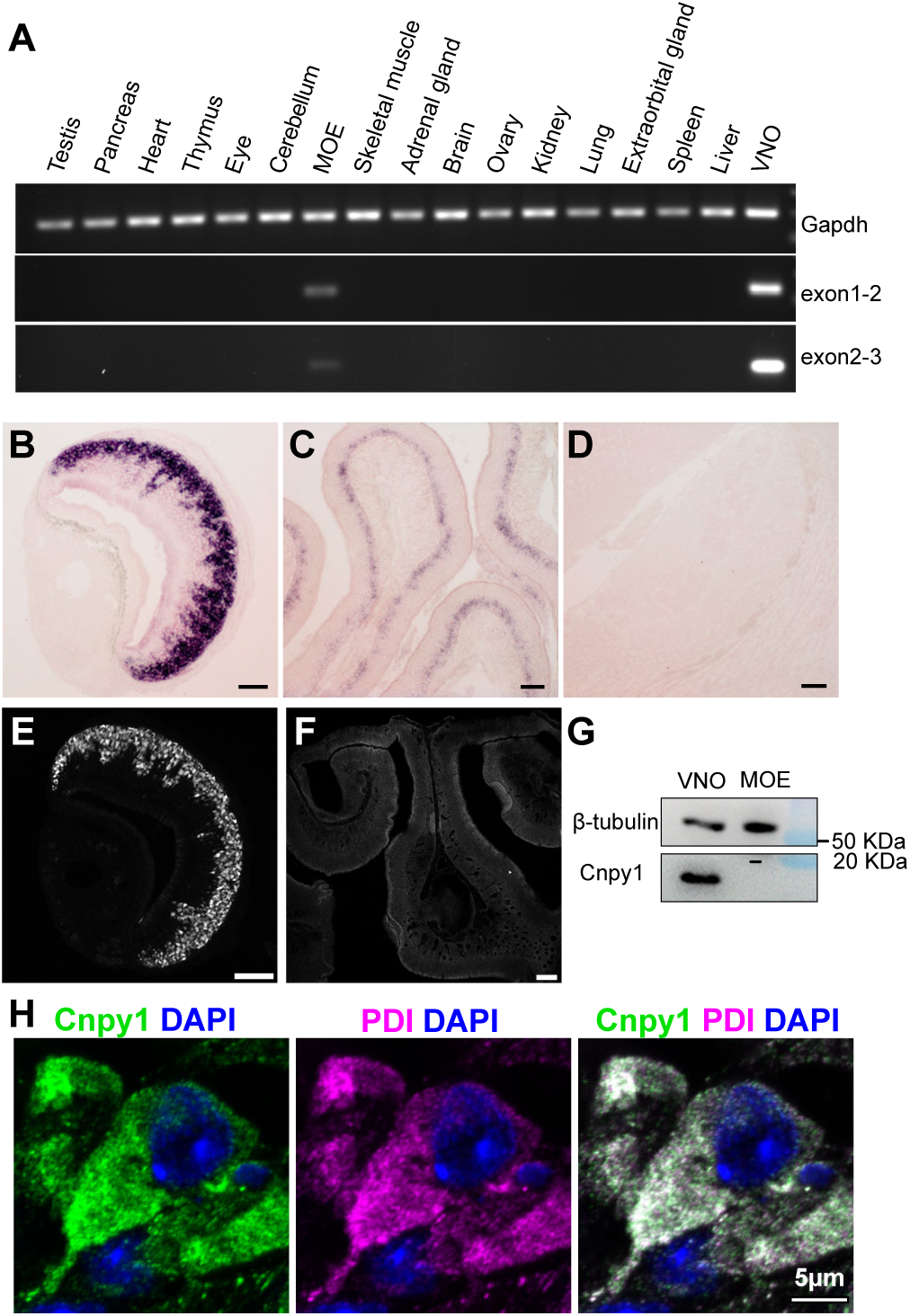
Cnpy1 is a Gαo VNO neuron specific ER protein. **A**) PCR amplification with primers specific to Cnpy1 exon1-2 (Cnpy1-FL) or exon 2-3 (common to Cnpy1-FL and Cnpy1-205) using cDNA from various mouse tissues as template. Amplicons visualized by agarose gel electrophoresis, show strong amplification in the vomeronasal organ (VNO) and weaker amplification in the main olfactory epithelium (MOE), with *Gapdh* serving as a control for cDNA integrity. **B-D**) RNA in-situ hybridization using anti-sense Cnpy1 riboprobe shows strong signal in VNO coronal sections (**B**), lower in MOE (**C**), and no expression in accessory olfactory bulb (**D**). **E, F**) Immunofluorescence using anti-Cnpy1 custom antibody shows protein expression specifically within basal VNO neurons (**E**), but not in MOE (**F**). **G**) Western blot shows Cnpy1 band in VNO but not MOE lysate, with β-tubulin as a loading control. **H**) High magnification image of VNO sensory neuron labelled with anti-Cnpy1 (green), anti-PDI (magenta) as endoplasmic reticulum (ER) marker, and DAPI (blue) marking the nucleus. Cnpy1 is co-localized in the ER. B-F scale bar is 100 μm.

Next, to test whether Cnpy1 protein follows the same pattern as its mRNA expression and to study its function, we generated a custom peptide antibody specific to Cnpy1. Immunolabeling of tissue sections revealed a localization pattern similar to Cnpy1 mRNA in the basal zone (**Figure 2E**). Interestingly, we did not see any Cnpy1 protein in the MOE (**Figure 2F**), even though small amount of transcript was detected, indicating the possibility of post-transcriptional regulation of Cnpy1 expression. Western blot analysis of Cnpy1 from VNO and MOE tissue lysates further validated this result, showing a robust band in VNO but absence in MOE (**Figure 2G**). From our single cell transcriptomics data (**Figure 1A, 1B**) and from RNA-ISH (**Figure 2B**) *Cnpy1* expression appeared to be restricted to basal/ Gαo neurons. To confirm if Cnpy1 is exclusive to Gαo neurons, we immunolabeled VNO sections using anti-Cnpy1 together with anti-G⍺o or anti-Nsg1 (marker for Gαi2 neurons) along with the antibody against ER retention sequence - SEKDEL, that preferentially marks the cell bodies of Gαo neurons (**Figure S2A, S2B**), (15). This showed that Cnpy1 expression is restricted to Gαo neurons (**Figure S2A**) with no labeling in Gαi2 neurons (**Figure S2B**).

Cnpy homologs have been shown to be resident in the ER and function as chaperones of various membrane receptors such as Toll-like receptors and immunoglobulins (16, 17, 21–26). We hypothesized that Cnpy1 is also an ER resident protein and a putative ER chaperone. To identify the subcellular localization of Cnpy1, we co-labeled VNO sections with anti-Cnpy1 and anti-Protein Disulphide Isomerase (PDI), a known ER resident chaperone. High resolution confocal microscopy showed that Cnpy1 is co-localized with PDI in the ER of Gαo neurons (**Figure 2H**), indicating that similar to other Cnpy homologs, mouse Cnpy1 is also likely to function in the ER of Gαo VSNs. Further, we cloned Cnpy1 with a N-terminal myc tag after the start codon, or C-terminal myc before its stop codon, expressed the tagged Cnpy1 proteins via transient transfection in HEK293 cells followed by detection of the expressed protein using anti-Cnpy1 or anti-myc tag antibodies. myc-Cnpy1 could only be detected with anti-Cnpy1, but not with anti-myc (**Figure S3A**), indicating that the N-terminal tag is cleaved off as a result of signal peptide cleavage, whereas C-terminal tagged (Cnpy1-myc) could be detected with both anti-Cnpy1 and anti-myc (**Figure S3B**). Cnpy1 was also observed to be localized to the ER in HEK293 cells (**Figure S3C**), similar to earlier studies that reported the localization of cloned Cnpy2-Cnpy5 to the ER in mammalian cell lines. Interestingly, mouse Cnpy1 C-terminal lacks a KDEL-type ER retrieval sequence, indicating that its ER retention may involve some other motif or mechanism.

### Loss of Cnpy1 leads to reduction of V2R expressing Gαo neurons

After confirming that Cnpy1 is an ER protein, we took a loss of function approach to identify the role of Cnpy1 in Gαo neurons. Crispr-Cas9 mediated targeting resulted in a 99 base pair deletion leading to a non-functional form of Cnpy1 protein to generate a Cnpy1-knockout (Cnpy1^-/-^) mouse model (**Figure 3A**). Cnpy1^-/-^ mice did not show any developmental abnormalities, nor any obvious breeding problems and could be maintained as a homozygous knockout line. VNOs collected from Cnpy1^-/-^ mice showed a complete absence of Cnpy1 protein, as characterized by western blot from lysate and immunolabeling of VNO sections (**Figure 3B, 3C**). To identify the effect of loss of Cnpy1, we performed RNA sequencing (RNAseq) of VNO from three replicates each of Cnpy1^+/+^ and Cnpy1^-/-^ littermate mice, at postnatal day 60 age (P60). We identified 111 genes differentially expressed between Cnpy1^-/-^ and Cnpy1^+/+^ with adjusted p-value less than 0.01 and absolute fold change of 1.5. Of these, 84 genes are downregulated and 27 are upregulated in Cnpy1^-/-^ compared to Cnpy1^+/+^ (**Figure 3D, S4A, Dataset S1**). Among downregulated genes, the major group that stood out is type-2 vomeronasal receptor genes (V2Rs) with about 66 V2R genes. This constitutes almost 50% of total V2Rs annotated from the mouse genome. To validate the RNAseq results, we performed qPCR using gene specific TaqMan probes against some of the V2Rs (Vmn2r65, 54, 53, 24, 112, 110, 116, 42, 2r1) and observed their downregulation in Cnpy1^-/-^ compared to Cnpy1^+/+^ VNO (**Figure S4B**). Of the 6 V2Rs observed to be upregulated in RNAseq, Vmn2r72 upregulation was also validated by qPCR assay (**Figure S4B**). Based on their sequence homology, V2Rs are classified into families A, B, C, D. Members of the A,B,D families largely follow the one-neuron-one-receptor rule, whereas family-C V2Rs are broadly expressed across Gαo neurons, where they are co-expressed with an ABD V2R per cell (27–29). We observed that downregulated V2Rs from **Figure 3D** were dispersed across the family tree, with representation across all families (**Figure S5**). These data indicate that the loss of Cnpy1 significantly impacts specific V2Rs across all families, as seen at the bulk RNA level.

**Figure 3.**
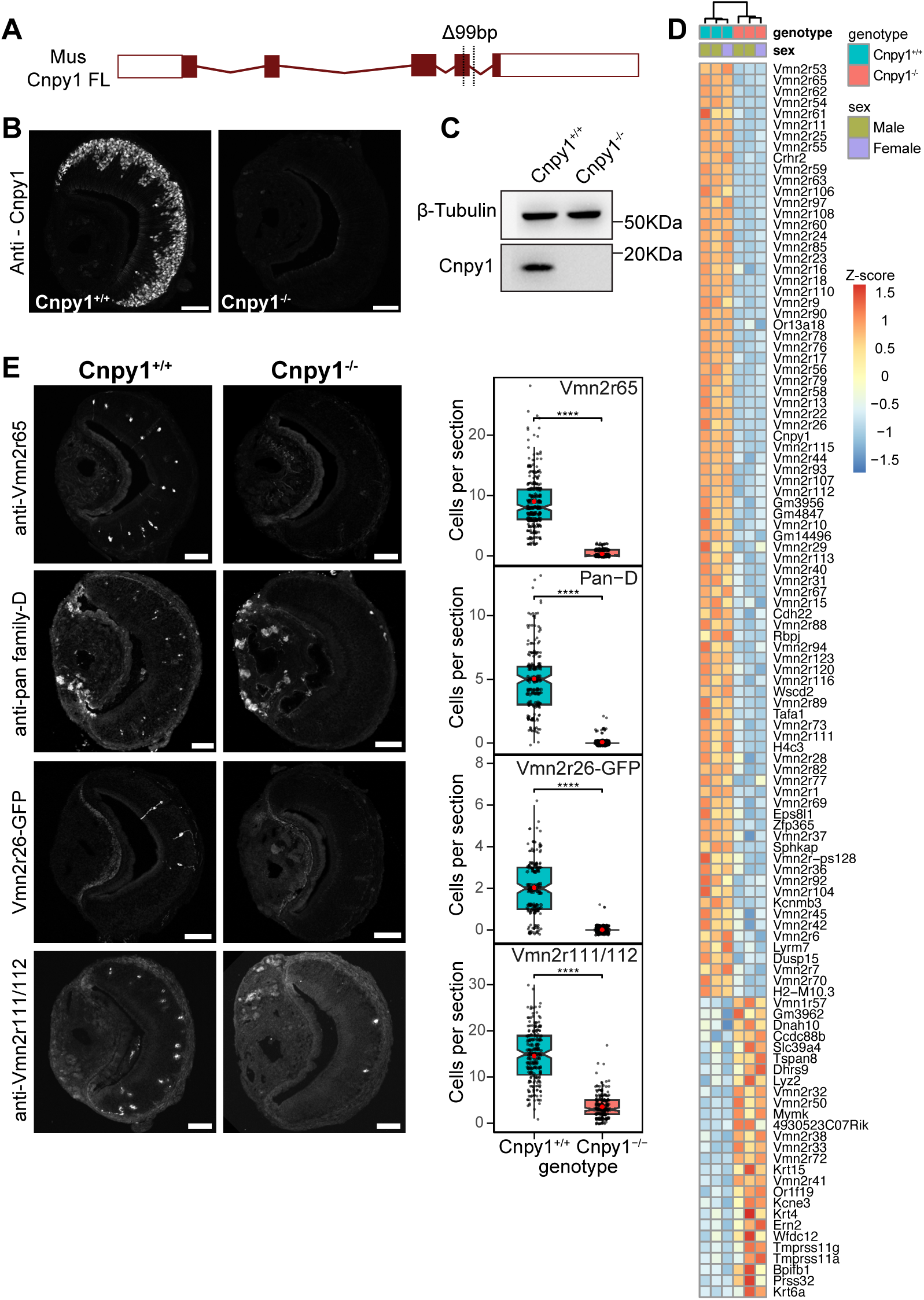
Deletion of Cnpy1 results in downregulation of multiple V2Rs **A)** Schematic of Cnpy1 gene showing a 99 base pair deletion in exon-4 leading to generation of a Cnpy1^-/-^ mouse line. **B, C**) Absence of Cnpy1 protein confirmed by anti-Cnpy1 immunofluorescence labeling of VNO section (**B**) and west-ern blot (**C**), with β-tubulin as loading control. **D**) Heatmap of differentially expressed genes from bulk RNA sequencing performed on Cnpy1^-/-^, Cnpy1^+/+^ littermate mice (3 each) VNOs. 111 genes with an adjusted p-value < 0.01 and an absolute fold change of 1.5 are shown ordered by fold change (most downregulated genes on the top). Of the 84 downregulated genes, 66 code for V2R (Vmn2r) GPCRs. **E**) Immunolabeling of P60 VNO sections with antibodies against Vmn2r65, pan-family D V2Rs, Vmn2r111/112, and GFP neurons representing Vmn2r26 shows absence of these V2R expressing neurons in Cnpy1^-/-^ VNOs compared to Cnpy1^+/+^. Representative images are on the left; box plots on the right show quantification of cells per section from at least 50 sections for each of 3 replicates per genotype. The notch on the box plots indicates the median, and red dot denotes mean. Statistical significance was assessed using Mann–Whitney U test. **** indicates p value < 10-4. Scale bar: 100 μm.

Next, we sought to investigate V2R downregulation at the protein level in individual neurons using immunohistochemistry. Using custom generated anti-Vmn2r65, anti-Vmn2r111/112 antibodies, and anti-pan family-D (Vmn2r53-56) (30), we performed immunolabeling on VNO sections from P60 Cnpy1^-/-^, Cnpy1^+/+^ littermate mice and counted the number of neurons expressing that V2R per section. We also crossed a mouse line that co-expresses GFP in Vmn2r26 expressing neurons (*Vmn2r26-IRES-tauGFP*, also termed V2r1b-IRES-tauGFP) (31), with Cnpy1^-/-^ to count Vmn2r26 neurons in the absence of Cnpy1. These V2Rs (Vmn2r65, family-D, Vmn2r26-GFP, Vmn2r111/112) representing multiple families, showed almost no labeling in Cnpy1^-/-^compared to Cnpy1^+/+^ VNOs (**Figure 3E, S6**), indicating the complete absence of neurons expressing these V2Rs. Lack of V2R expressing cells in Cnpy1^-/-^ indicates that the reduction of receptor mRNA levels seen with bulk RNA seq is a consequence of reduced cell numbers expressing those receptors rather than attenuated expression at an individual neuron level.

We asked whether the absence of multiple V2R expressing neurons in Cnpy1^-/-^ VNO results in a reduction in the total number of Gαo neurons. To address this question, we immunolabeled VNO sections with anti-Nsg1 and anti-Gαo antibodies, markers of the apical/basal zone (**Figure S2A**) VSNs respectively. Quantification of the demarcated area in at least 50 sections each from 3 replicates of P60 Cnpy1^-/-^ and Cnpy1^+/+^ littermate mouse VNOs revealed a 35.4% reduction in mean Gαo/basal zone area (**Figure 4A**). Concomitant with a reduction of Gαo neurons, the total sensory epithelium area also showed a reduction in Cnpy1^-/-^ mouse VNOs.

**Figure 4.**
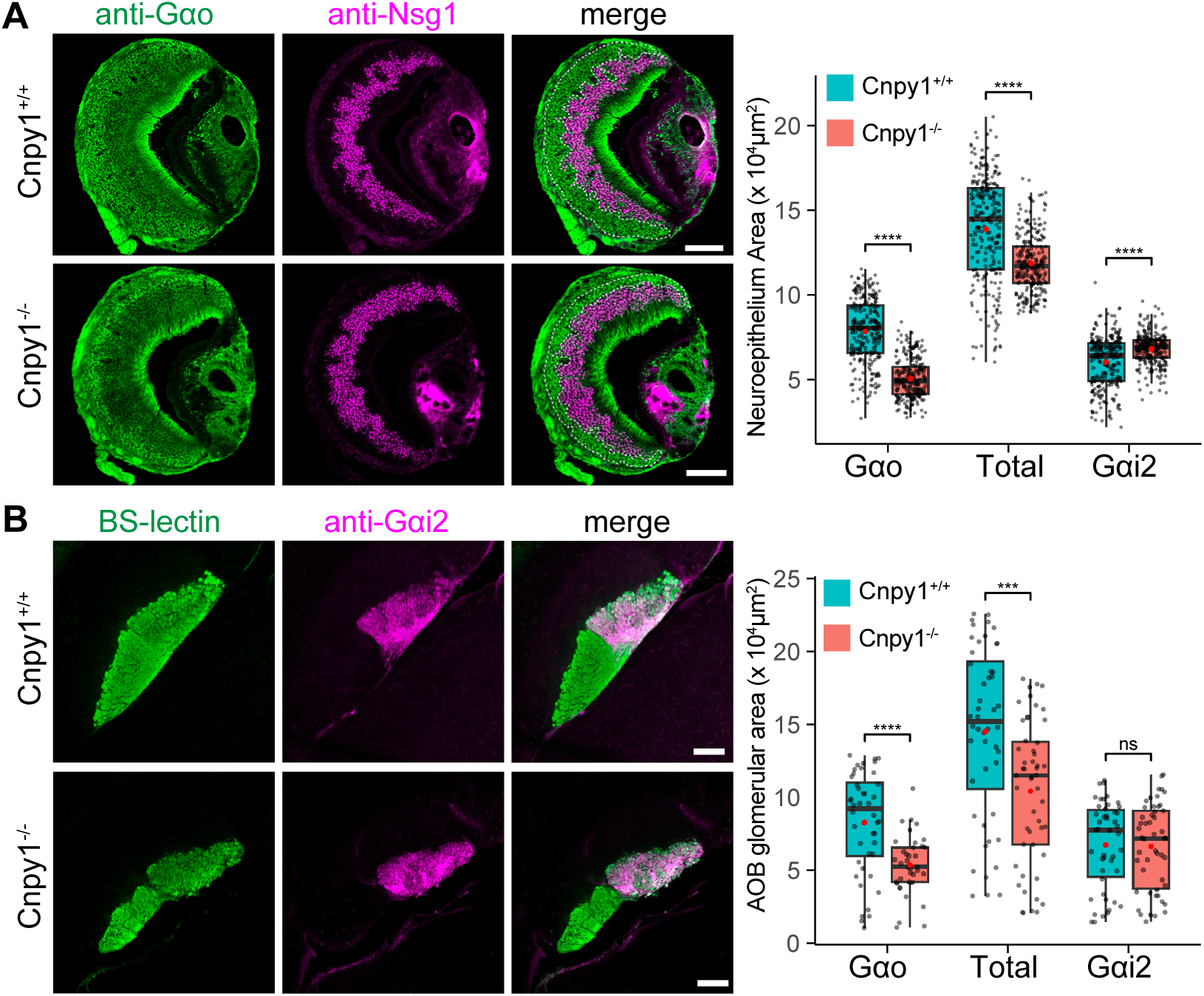
Gαo zone is reduced in Cnpy1^-/-^ mice. **A)** VNO cryosections from P60 Cnpy1^-/-^ and Cnpy1^+/+^ mice immunolabelled to mark Gαo or Gαi2 zones (anti-Nsg1/Neep21). Representative images (left) show the area (dotted line) that was quantified (right) from 3 pairs of littermate mice (Cnpy1^+/+^:244, Cnpy1^-/-^: 238 sections). **B**) Representative images (left) of AOB vibratome sections co-labelled with BS-lectin that marks the combined glomerular and nerve layer area along with anti-Gαi2, followed by their quantification (3 animals, Cnpy1^+/+^: 44, Cnpy1^-/-^: 37 sections) on the right. Gαo area was quantified by subtracting Gαi2 from the total area. Both VNO and AOB quantifications show a substantial reduction in Gαo area, which results in a thinning of the total neuroepithelium. There is no reduction in Gαi2 area. Horizontal line on the box plots indicates the median and red dot denotes mean. Statistical significance was assessed using Mann–Whit-ney U test. **** - p value < 10-4; *** - p value < 10-3; ns - not significant. Scale bar: 100 μm.

Apical or basal VSNs project axons to respective segregated glomerular regions of the AOB. Quantification of the glomerular region area from sagittal sections of the AOB marked by Banderiraea Simplicifolia (BS)-Lectin (32) and negatively labeled by anti-G⍺i2 also revealed a specific reduction in the basal zone region (**Figure 4B**). These data demonstrate that in the absence of Cnpy1, multiple V2R expressing neurons are lost, resulting in a substantial reduction in the Gαo neuroepithelium zone. Thus, the downregulation of almost 50% V2R genes seen in Cnpy1^-/-^ RNAseq data (**Figure 3D**) could be attributed to the loss of these neurons.

### Loss of Cnpy1 leads to death of specific Gαo VSNs

Since we observed a substantial loss of V2R expressing Gαo VSNs in P60 Cnpy1^-/-^ mice, we asked whether these neurons were missing at birth of the animal or are lost later during postnatal development. We counted the numbers of V2R labeled VSNs, (shown in **Figure 3E, S6**) in sections from mice with and without Cnpy1 at different ages. At age P7, V2R expressing neurons were present in Cnpy1^-/-^ and their numbers comparable with Cnpy1^+/+^. However, by P15 there was a clear reduction of these V2R expressing neurons in Cnpy1^-/-^ VNOs that persisted into later ages (**Figure 5A-C, S7A**). This indicates that VSN numbers are normal at birth and decrease during postnatal development. To understand how neurons are being lost, we labelled VNO sections from Cnpy1^-/-^ and Cnpy1^+/+^ mice across postnatal ages with an antibody against cleaved caspase 3 (CC3), a marker for apoptotic neurons (33) along with anti-SEKDEL, a marker for Gαo neurons (**Figure 5D**). This allowed us to quantify CC3 positive cells within Gαo zone and entire neuroepithelium at each postnatal age. The total number of CC3+ cells showed a significant increase in P15 Cnpy1^-/-^ VNOs compared to Cnpy1^+/+^ controls (**Figure 5E**). CC3+ cell numbers within Gαo zone (**Figure S7B**) and normalizing these to a declining postnatal Gαo zone in Cnpy1^-/-^ VNOs (**Figure 5F**) show consistently elevated levels P15 onwards, compared to age matched Cnpy1^+/+^ VNOs (**Figure 5G**). Immunolabeling of P15 Cnpy1^-/-^ sections with anti-CC3 and anti-OMP, confirmed that the elevated number of apoptotic cells in Cnpy1^-/-^ Gαo zone were OMP labelled mature neurons (**Figure S7C**). Further, VNO sections from P60 Cnpy1^-/-^ co-labelled with anti-OMP, anti-Gαo confirmed that residual Gαo neurons in the neuroepithelium of Cnpy1^-/-^ mice were OMP positive, indicating that absence of Cnpy1 does not result in a developmental block prior to mature neuron stage (**Figure S7D**), as seen in the VNOs from mice lacking the transcription factor, ATF5 (34). Collectively, these data show that V2R expressing mature VSNs are lost due to apoptotic cell death during early postnatal development after Gαo fate specification in Cnpy1^-/-^ VNO.

**Figure 5.**
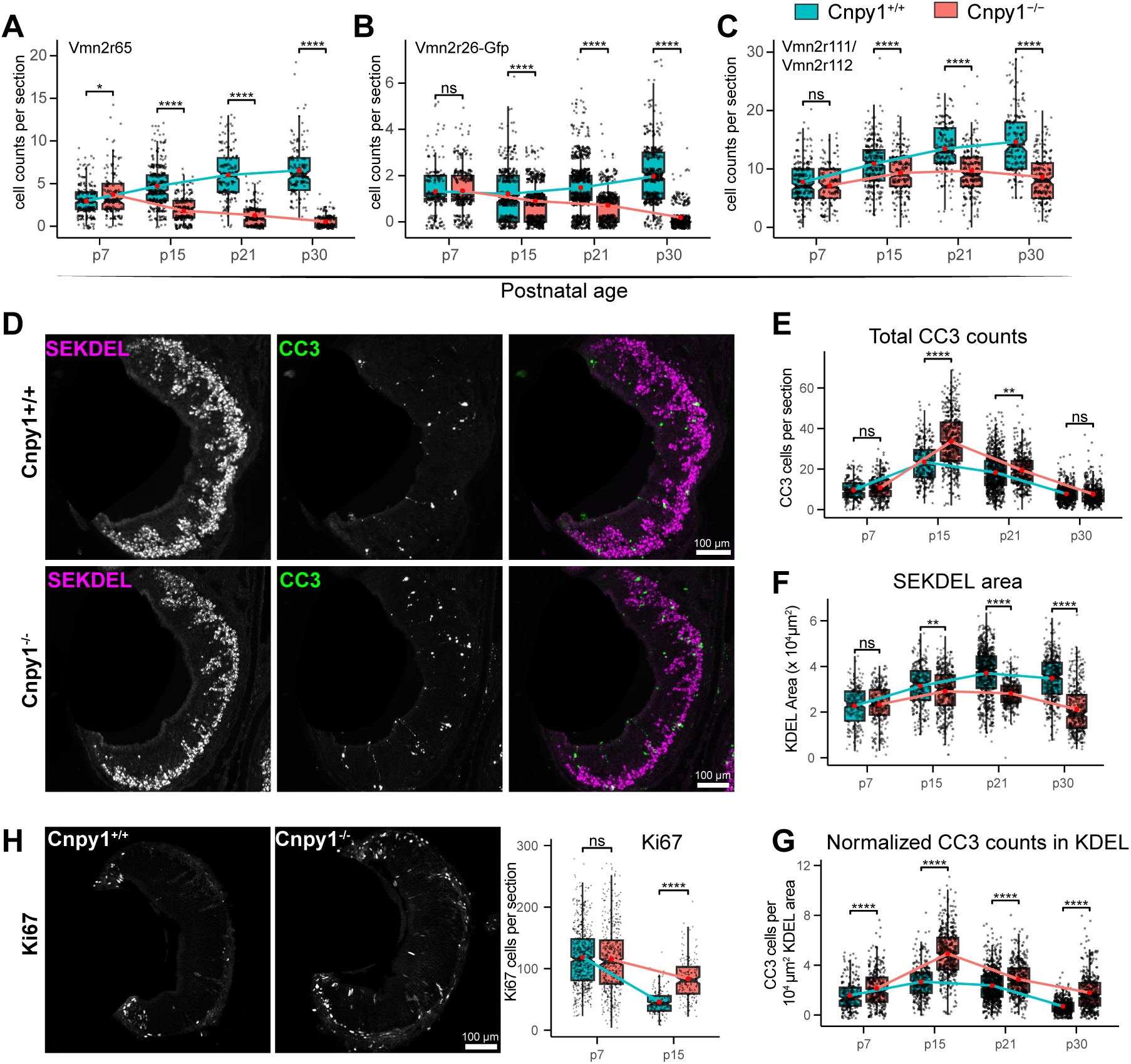
Enhanced cell death and neurogenesis in Cnpy1^-/-^ VNO neuroepithelium. **A-C**) Quantification of V2R expressing neurons from Cnpy1^-/-^ and Cnpy1^+/+^ VNO cryosections across postnatal ages, using anti-V2R antibodies shown in Figure 3. Counts of Vmn2r65 (**A**), Vmn2r26 (**B**), Vmn2r111/112 (**C**) are comparable at P7. In Cnpy1^+/+^ number of neurons expressing these V2Rs increase with postnatal development, but in Cnpy1^-/-^ VNOs, they decline from P15 onwards. **D-G**) Representative images of P15 VNO cryosections labelled with anti-SEKDEL to mark Gαo zone, and Cleaved Caspase-3 (CC3), a marker of apoptotic cells. CC3 positive cells quantified in the neuroepithelium (**E**) show an increase in P15 Cnpy1^-/-^VNO. The Gαo zone (SEKDEL) area in Cnpy1^-/-^ and Cnpy1^+/+^ VNOs is comparable at P7 but declines from P15 onwards (**F**). The total number of CC3 cells within Gαo area, as well as CC3 numbers normalized to the declining Gαo area at respective age groups (**G**), show elevated Gαo zone specific apoptosis across postna-tal ages, with a peak at P15 in Cnpy1^-/-^. **H**) Labeling and quantification of VNO sections with anti-Ki67 indicates elevated neurogenesis at P15 age in Cnpy1^-/-^ VNO compared to Cnpy1^+/+^. All quantifications were performed with 5 animals per age and genotype and at least 150 sections per condition. Box plot notches indicate the median and red dot denotes mean. Statistical significance was assessed using Mann–Whitney U test. **** - pvalue < 10-4; *** - p value < 10-3; ** p value < 10-2; * - p value < 0.05; ns - not significant.

One of the hallmarks of olfactory and vomeronasal neurons, is their continuous regeneration from a stem cell population, which in case of VSNs leads to a developmental transition and cell fate bifurcation via an immature Gαi2 or Gαo state to respective mature neurons (35). To address whether there is a change in the rate of neurogenesis triggered due to loss of Gαo neurons, we labelled VNO sections for Ki67, a marker of cell proliferation (36). Quantification of Ki67 positive cells revealed a marked increase in P15 Cnpy1^-/-^ mouse VNOs (**Figure 5H**), indicating a consistent and compensatory neurogenesis accompanying the VSN loss due to cell death. Collectively these data indicate that the absence of Cnpy1 results in a loss of V2R neurons due to cell death during postnatal development that is likely to be sensed and compensated to some extent by enhanced neurogenesis.

### Cnpy1 interacts with V2Rs and other ER chaperones

Our next step was to understand the molecular function of Cnpy1 in Gαo neurons. Cnpy1 localization to the ER, the function of homologous Cnpy proteins as ER chaperones, and the loss of several V2R expressing VSNs in Cnpy1^-/-^ mice, lead us to ask whether Cnpy1 directly interacts with V2Rs, acting as a specific chaperone. To test this, we immunoprecipitated Cnpy1 from lysate of C57BL/6J (B6) VNOs using anti-Cnpy1 antibody (IP_Cnpy1:B6_). As controls, we performed IP from Cnpy1^-/-^ VNO lysate (IP_Cnpy1:KO_) as well as an IP using normal IgG from B6 VNO lysate (IP_IgG:B6_). The protein eluates from all IP samples were fragmented to peptides using trypsin and labeled with TMT. We pooled TMT labelled samples and performed fractionation followed by liquid chromatography-quantitative mass spectrometry to identify the co-immunoprecipitating interacting partners of Cnpy1. Proteins in IP_Cnpy1:KO_ and IP_IgG:B6_ serve as negative controls to identify non-specific interactions. Therefore, we considered common enriched hits based on absolute foldchange more than 2, calculated from [IP_Cnpy1:B6_ vs IP_Cnpy1:KO_] and [IP_Cnpy1:B6_ vs IP_IgG:B6_] as true interacting partners of Cnpy1. With this process, we identified 27 specific interacting partners: 13 V2Rs, 5 ER chaperones and 9 other proteins of various functions (**Figure 6**, **Dataset 2**). Due to high sequence homology, the identification of specific V2Rs via MS is likely to be an underestimate, because only those trypsin cleaved peptides that can be uniquely mapped to individual V2R sequences result in the identification of interacting Vmn2r members. The interaction of Cnpy1 with V2Rs and other ER chaperones indicates that it is a putative V2R specific chaperone that might function along with other chaperones for the proper folding of V2Rs in the ER of Gαo VSNs.

**Figure 6.**
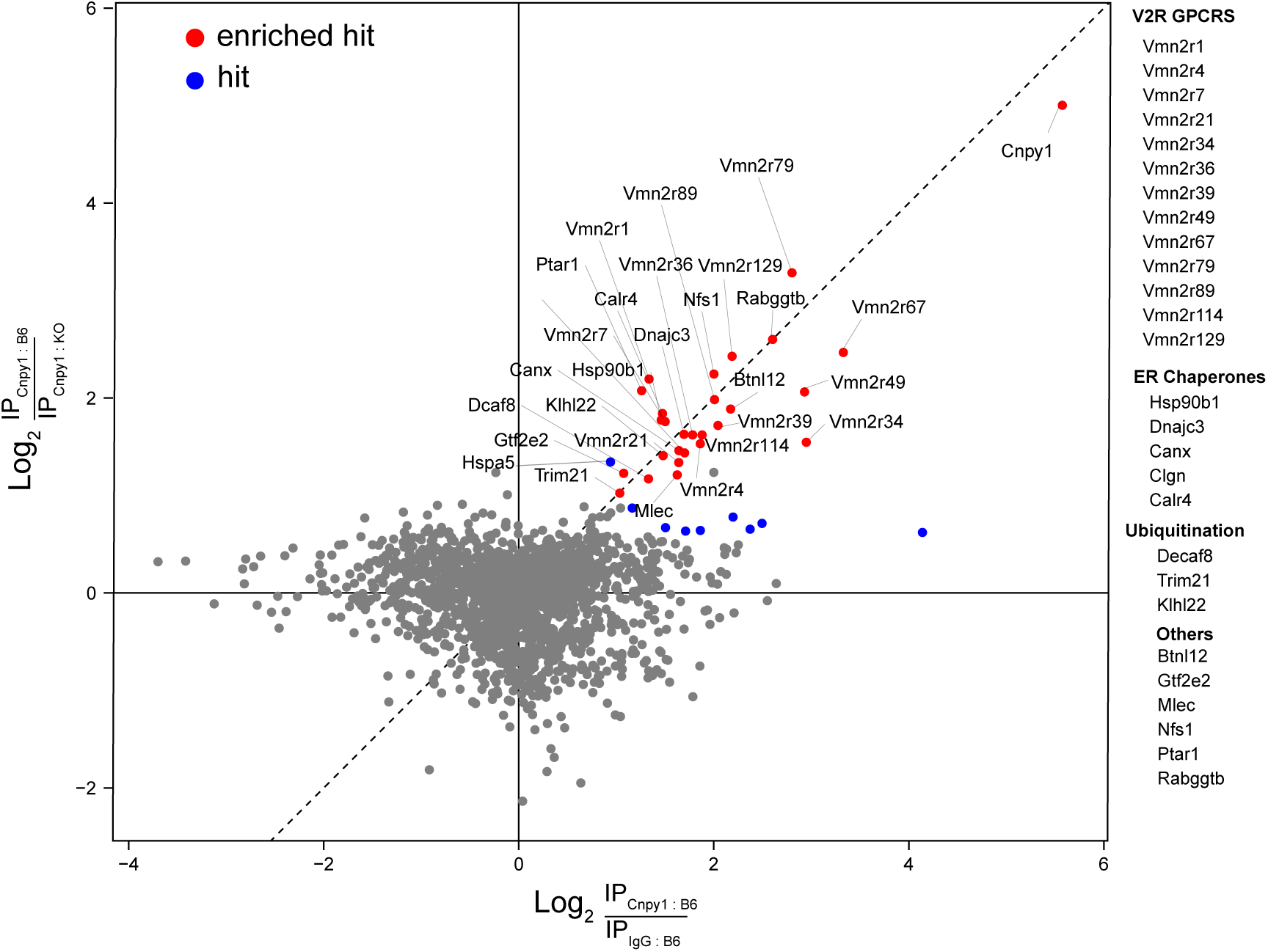
Cnpy1 interacts with Vmn2r GPCRs and ER chaperones. Immunoprecipitation (IP) of Cnpy1 followed by quantitative mass spectrometry to identify specific interacting partners of Cnpy1. Cnpy1 IP performed using anti-Cnpy1 from C56BL/6 (B6) mouse VNO lysates (IP_Cnpy1:B6_) was compared with two negative controls: normal rabbit IgG IP from B6 VNO lysate (IP_IgG:B6_) and anti-Cnpy1 IP from Cnpy1^-/-^ (knockout - KO) VNO lysate (IP_Cnpy1:KO_). Each IP was performed in 3 biological replicates, each replicate containing pooled VNOs from 50 mice, except for IP_IgG:B6_ which was performed in duplicate. Specific hits are identified by comparing fold change of signal intensity from [IP_Cnpy1:B6_ over IP_Cnpy1:KO_] (Y axis) and [IP_Cnpy1:B6_ over IP_IgG:B6_] on X axis. Proteins with an absolute fold change ≥ 2 and a false discovery rate (FDR) < 0.05 in both comparisons and falling along the diagonal are considered specific interacting proteins (enriched hits) and are shown in red. A list of enriched hits is provided on the right. Hits with FDR < 0.05, showing an absolute fold change ≥ 2 in one comparison and between 1.5 and 2 in the other, are highlighted in blue.

### Cnpy1 loss does not activate the canonical unfolded protein response or disrupt V2R trafficking and glomerular targeting

To investigate whether V2Rs are exported out of the ER of VSNs in the absence of Cnpy1, we imaged immunolabeled Vmn2r65, Vmn2r111/112 and Vmn2r2 towards the lumen of the vomeronasal sensory epithelium, where VSNs project their dendritic tips. Since VSNs expressing these V2Rs are mostly lost by P30 age in Cnpy1^-/-^ mice, we used VNO sections from 2 weeks old mice, where these VSNs are still detected in Cnpy1^-/-^, although with reduced numbers (**Figure 5A**). We observed an equivalent amount of signal in Cnpy1^-/-^ as well as Cnpy1^+/+^ dendritic tips for immunolabeled V2Rs (**Figure 7A, 7B, S8**), indicating that post-ER export of V2Rs along the secretory pathway and their targeting is unlikely to be defective due to the absence of Cnpy1. Furthermore, we reasoned that if there was a build-up of misfolded V2R GPCRs in Cnpy1^-/-^ VSNs, it should result in an unfolded protein response (UPR). Using a combination of approaches, we probed P14 VNOs for molecular mediators representing the three major arms of the UPR pathway (37, 38). Splicing of the X-box binding protein-1 (*Xbp1*-s) transcription factor mRNA -probed with regular and quantitative PCR (**Figure 7C, 7D**); levels of phosphorylated Eif2ak3/PERK and eukaryotic translation initiation factor 2A (phospho-PERK, phospho-Eif2a) probed by western blot (**Figure 7E, 7F**); the transcription factor Ddit3/CHOP -probed by TaqMan q-PCR (**Figure 7D**), as well as ATF5 levels probed by western blot and immuno-fluorescence (**Figure 7G-I, S9**), show comparable levels between Cnpy1^-/-^ and normal mouse VNOs. Of note, Gαo VSNs normally exhibit high ER stress-like condition -as demonstrated in their elevated levels of ER chaperones, hypertrophic ER membrane (15) - and in the levels of p-PERK, p-Eif2a shown here (**Figure 7E, 7F**), which seem to resemble those seen in tunicamycin treated mouse liver (used as a positive control for ER stress). Nevertheless, the deletion of Cnpy1 does not seem to alter this higher baseline level ER stress-like state (**Figure S10**) in VNO neurons.

**Figure 7.**
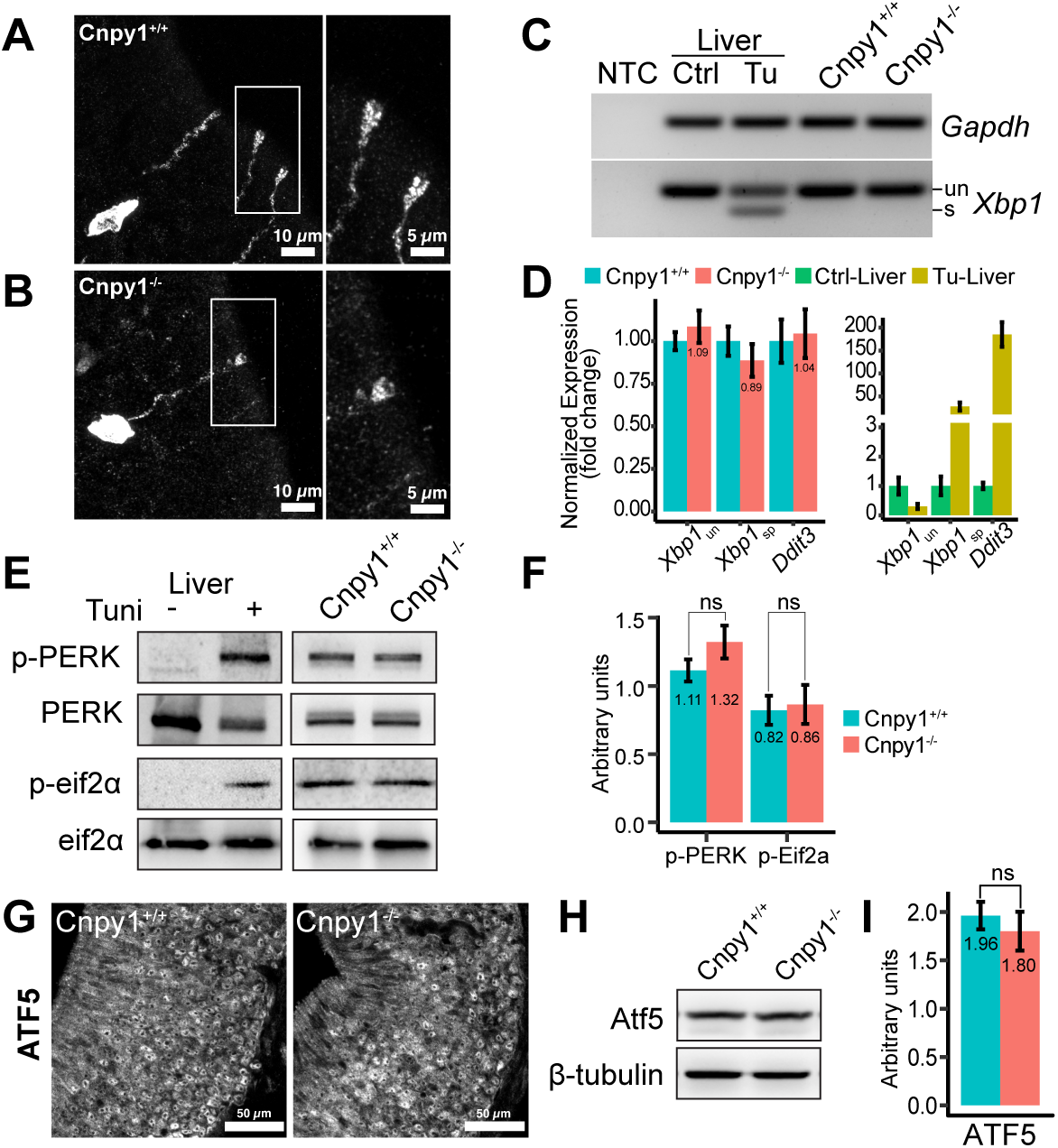
Normal trafficking of V2R GPCRs and lack of canonical UPR activation in Cnpy1-null VSNs. **A, B**) Immunofluorescence confocal images of anti-Vmn2r65 labeling on P15 Cnpy1^+/+^ and Cnpy1^-/-^ VNO sections, show comparable GPCR localization to VSN cell bodies and dendritic tips in both genotypes. Additional images of Vmn2r65, Vmn2r2, Vmn2r111/112 localization to microvilli at the sensory lumen, are shown in **Figure S8**. **C**) *Xbp1* PCR-gel electrophoresis detecting spliced (s) and unspliced (un) bands. *Xbp1*s-indicative of active ER stress/unfolded protein response (UPR) is detected in tunicamycin treated mouse liver cDNA (Tu; positive control), but absent from Cnpy1^+/+^ and Cnpy1^-/-^ P14 VNO cDNA. *Gapdh* was used a control for cDNA integrity. NTC, no template control. **D**) Quantification of *Xbp1*s and Ddit3/CHOP levels from cDNA by qPCR. Xbp1s, Ddit3 levels remain comparable between genotypes, whereas both show increase in tunicamycin-treated liver (Tu-Liver) relative to vehicle control, confirming assay sensitivity. **E, F**) Western blots of phosphorylated Eif2ak3/PERK (p-PERK) and phosphorylated Eif2α (p-Eif2α) confirm UPR activation in Tu-liver lysates (positive control). p-PERK and p-Eif2α are detectable in P14 VNO lysates; however, band intensities and their quantification normalized to β-tubulin loading controls reveal comparable levels between Cnpy1 genotypes. **G–I)** ATF5 levels are similar across Cnpy1 genotypes in P14 VNO sections and lysates, as assessed by immunofluorescence and immunoblotting. Immunoblot quantification normalized to β-tubulin is shown in **I**. Quantification bar plots are represented as mean ± SEM of at least 3 biological replicates. ns, not significant (P > 0.05, Welch’s t-test).

Since V2R GPCRs are known to be involved in the process of targeting axonal projections to the AOB (39, 40), we first investigated whether V2R GPCRs could be detected in the AOB, where Gαo VSNs project their axons to form glomeruli and synapse with mitral cells. Immunolabeling for multiple V2Rs failed to detect their presence in normal as well as Cnpy1^-/-^ AOB sections at P14 (**Figure S11**). This is consistent with previous reports where robust V2R labelling is detected in the VNO, but not the AOB (41) and indicate that V2Rs may play a permissive but not instructive role for axonal guidance to the AOB. We next investigated glomerular targeting in Vmn2r26-GFP;Cnpy1^-/-^ P7 mice, when Vmn2r26-GFP neurons can still be detected. GFP labeled axons still coalesce towards the posterior AOB (**Figure S12**). This indicates that during early postnatal age, the gross anatomical wiring of V2R expressing VSNs to the AOB is not altered in Cnpy1^-/-^, although finer defects in glomerular or synapse formation may exist. Collectively, the data presented here rule out defective V2R protein export from the ER and UPR activation as a mechanism for cell death in Cnpy1^-/-^ Gαo VSNs.

### Reduced neuronal activation and behavioral defects in Cnpy1-/- mice

We next investigated whether Gαo VSN activation, downstream of V2R GPCR signaling was affected. Cnpy1^-/-^ and Cnpy1^+/+^ P60 littermate mice were exposed to rat bedding, a source of predator kairomones or opposite sex bedding as a source of pheromones. Both stimuli were shown to be complex sources of ligands for VSNs, and their activation can be measured by immunolabeling VNO sections with an antibody to phosphorylated S6 ribosome subunit (pS6) (42). We immunolabeled VNO sections from mice exposed to these stimuli with anti-pS6 and anti-SEKDEL (to mark Gαo zone) and quantified the number of pS6 labeled neurons per section (**Figure 8A-8D**). In case of both stimuli, Cnpy1^-/-^ VNOs showed a clear reduction in the number of activated neurons within Gαo zone compared to Cnpy1^+/+^. These data indicate that despite loss of Gαo neurons in P60 Cnpy1^-/-^ mice due to cell death, the persisting neurons, show significantly reduced activation by ligands. Interestingly, there was a small but significant increase in pS6 cells outside the Gαo zone, which matches with a slight increase in Gαi2 zone we observed earlier (**Figure 4A**), pointing to the possibility of a compensatory increase in the number of Gαi2 neurons in Cnpy1^-/-^ neuroepithelium.

**Figure 8.**
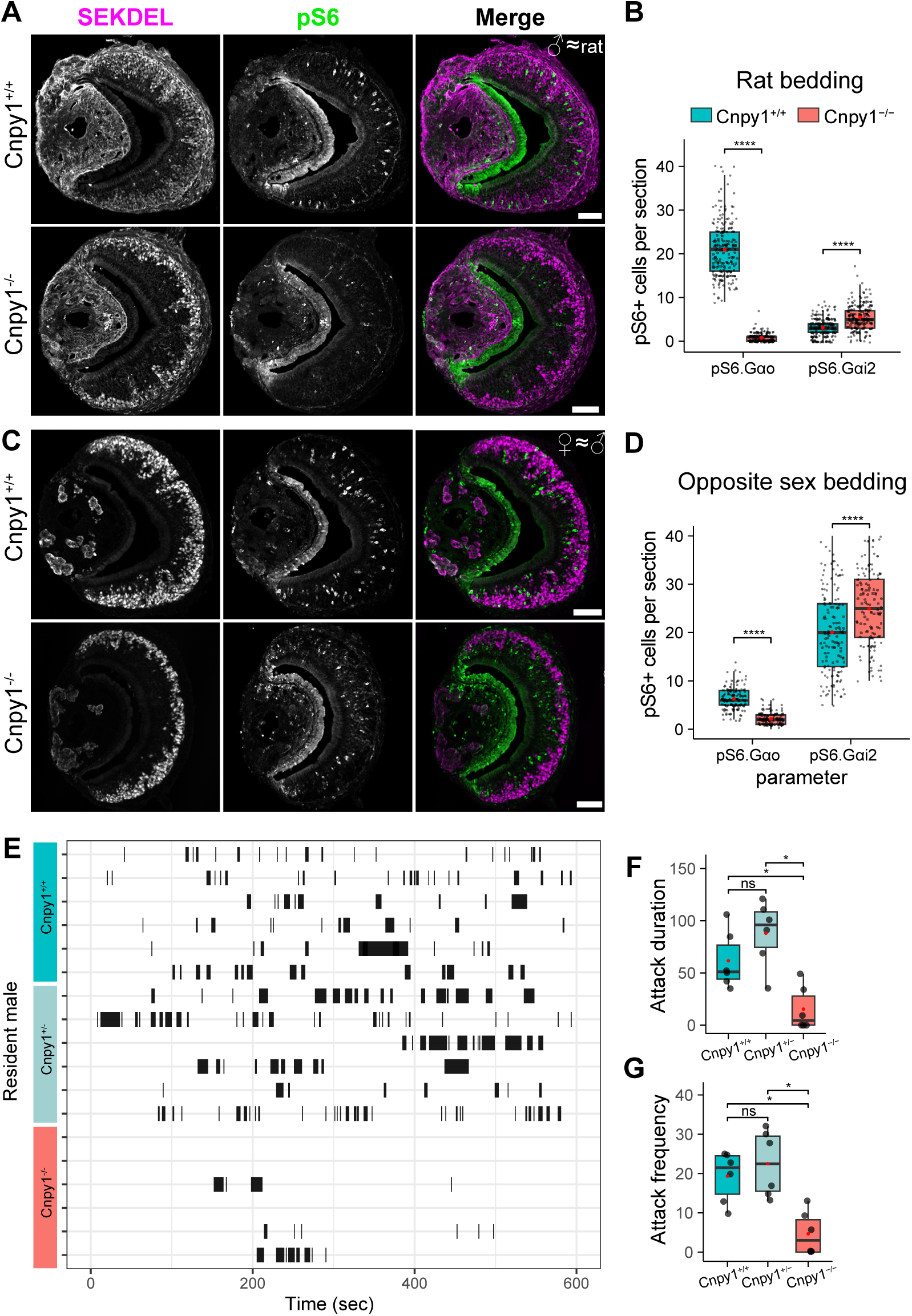
Impaired Gαo neuron activation and behavior in Cnpy1^-/-^ mice. **A-D**) Representative images and quantification of activated neurons in VNO sections from P60 Cnpy1^-/-^ and Cnpy1^+/+^ mice after exposure to rat bedding (predator signal; **A, B**) or opposite sex bedding (**C, D**). Activated VSNs are detected by anti-pS6 (green), and sections are co-labelled with anti-SEKDEL (magenta) to mark Gαo zone neurons. Quantification of pS6+ cells from 3 biological replicates and at least 50 sections per mouse (**B, D**), shows substantially reduced activation of Cnpy1^-/-^ Gαo neurons compared to Cnpy1^+/+^. A small but significant increase in activated neurons outside Gαo zone (Gαi2) was also detect-ed. **E-G**) Raster plot showing attack events by a male resident mouse towards intruder mouse over time (**E**). Attack duration (**F**) and number of attacks (**G**) from six resident animals each from Cnpy1^+/+^, Cnpy1^+/-^ and Cnpy1^-/-^ are shown as box plots; the red dot denotes the mean. Cnpy1^-/-^ males exhibit markedly fewer attacks than mice having an intact Cnpy1 allele. Statistical significance was assessed using Mann–Whitney U test. **** - p value < 10-4; * - p value < 0.05; ns - not significant. Scale bar: 100 μm.

One of the behavioral consequences of loss of Gαo VSN activation is reduced male-male aggression in a classical resident-intruder paradigm (43). Hence, we exposed littermate resident male mice at P60 age to castrated male mice swabbed with male mouse urine. Cnpy1^+/+^ and Cnpy1^+/-^ mice reacted with aggressive behaviors towards the intruder male, while Cnpy1^-/-^ males showed markedly reduced aggressive behaviors towards intruders (**Figure 8E-G**). Collectively, these data indicate that loss of Cnpy1 impairs activation of Gαo VSNs by chemical cues, producing deficits in behaviors normally mediated by this neuronal population.

## Discussion

In this study, we identify mouse Cnpy1 as a VNO specific protein, with its expression restricted to the Gαo subtype of sensory neurons. Compared to other Cnpy homologs, mammalian Cnpy1 has remained enigmatic because of an incomplete SAPLIP domain. The novel Cnpy1 transcript we identified in the mouse VNO, includes an additional upstream exon with a new translation start site that completes full-length Cnpy1 protein by contributing two additional cysteines and a signal peptide. Mapping trypsin digested peptides from our Cnpy1 IP-proteomics gave additional confidence to the predicted protein sequence (**Figure S13**). *Cnpy1* was first described and cloned from zebrafish, where its expression during embryonic development was shown at the mid-brain hind-brain boundary (MHB) (16). Subsequent studies in mouse also reported *Cnpy1* mRNA at the MHB (44). In contrast, we found *Cnpy1* mRNA in the VNO and MOE, but Cnpy1 protein only in the VNO. We propose that mouse *Cnpy1* is post-transcriptionally regulated, without a role for Cnpy1 protein in the MOE or MHB. The absence of developmental defects in our *Cnpy1* gene deletion mouse model supports this hypothesis. While the existence of an as-yet undescribed alternate transcript cannot be ruled out, our Cnpy1 knockout allele and anti-Cnpy1 antibody binding motif are likely to be common to all potential isoforms, arguing against such a possibility.

Our experiments reveal that Cnpy1 plays an essential, non-redundant role in Gαo VSNs. Intracellular localization of Cnpy1 in VNO sections as well as expression of the cloned gene in heterologous cells demonstrates an ER localization. IP-proteomics from the VNO identified Cnpy1 interactions with multiple V2R GPCRs as well as specific ER chaperones such as Hsp90b1, Dnajc3, Calnexin, Calr4. Mammalian Cnpy homologs, Cnpy2 - Cnpy5 are known to function as ER co-chaperones. Cnpy3, Cnpy4 (also termed PRAT4A, 4B), along with Hsp90 family proteins, co-chaperone the folding and exit of Toll-like receptors in the ER of immune cells (22, 23, 26, 45–47). Likewise, Cnpy5 (also termed ERp1/MZB1), interacting with Grp94 and BiP, is an essential ER chaperone for the folding and secretion of immunoglobulins in B cells (24, 25, 48, 49). These studies along with our results, would point to a mechanistic role for mouse Cnpy1 in co-chaperoning V2R GPCRs.

Intriguingly, despite the absence of Cnpy1, post-ER targeting of V2Rs to dendritic tips and activation of canonical UPR/ER-stress were not affected. Instead, Cnpy1^-/-^ Gαo neurons exhibited reduced responses to VNO stimuli. While mice lacking Cnpy1 were born with normal number of VSNs, Gαo neurons were progressively and selectively lost during postnatal development due to apoptosis. A similar pattern–normal development of the accessory olfactory system till birth, followed by gradual postnatal loss of VSNs– has been described in multiple mouse models lacking GPCR signaling components in VSNs. For example, deletion of the Gαo G-protein subunit, essential for V2R signal transduction, results in selective postnatal apoptosis of this neuronal subtype (43, 50). Neurons in mice engineered to render a V2R nonfunctional, by insertion of LacZ in the coding region (39), also exhibit gradual postnatal loss. Similar phenotypes have been described in mouse models lacking a functional V1R (40), the cognate Gαi2 subunit (51), or TRPC2, the major vomeronasal ion-channel downstream of GPCR signaling (52). Collectively these studies highlight that functional vomeronasal GPCR signaling is essential for postnatal neuronal survival. We therefore propose that Cnpy1 is required in the ER of Gαo VSNs to assemble signaling-competent V2R GPCRs, the absence of which triggers apoptosis.

V2Rs belong to the class-C GPCR family, that contain a large extracellular ligand-binding Venus fly-trap domain, in addition to 7-transmembrane domains. Like other class-C GPCRs, V2Rs are likely to function as obligate homo- or heterodimers. Assembly of functional V2Rs may require participation of multiple ER chaperones, some of which could be Gαo VSN subtype specific. Cnpy1 could influence local folding, conformational states, posttranslational modifications of V2Rs their dimerization, or interactions with G-proteins. Defects in any of these could disrupt signaling, while escaping ER quality control. Interestingly, Cnpy4 has recently been proposed to influence Hedgehog pathway signaling by modulating membrane lipid homeostasis, raising the possibility that SAPLIP proteins contribute to cellular signaling via lipid organization or availability (53, 54). Whether this mechanism is conserved across other Cnpy homologs remains to be determined.

Although we observed substantial loss of V2R-expressing neurons in P60 Cnpy1-null mice, some neurons persisted. Since Cnpy1 does not affect VSN developmental maturation, therefore due to persistent neurogenesis, Gαo neurons are expected to be present at any age. In addition, Gαo neurons may differ in their individual susceptibility to a loss of receptor signaling, depending on the V2R they express. In the MOE, receptor identity has been shown to determine ER stress levels across neurons (55). While ER stress markers are generally elevated in VNO neurons, sensory GPCR induced ER stress may operate through distinct non-canonical mechanisms in the VNO (56). For instance, Cnpy2 has been shown function in context specific manner during ER stress either by activating the UPR or by reducing stress (21, 57). An alternate possibility is that some of the Cnpy1^-/-^ VSNs that fail to form a signaling competent V2R, may switch transcription to a different V2R, thereby transiently delaying apoptosis. In summary, our findings establish mouse Cnpy1 as a novel VNO specific ER chaperone, required for the survival of Gαo neurons. In conjunction with other ER chaperones, Cnpy1 likely contributes to the assembly of signaling-competent V2R GPCR complexes, opening new avenues for dissecting molecular mechanisms of vomeronasal receptor function.

## Materials and Methods

### Supporting Information Materials and Methods

#### Animals

All experiments were carried out with approval from the Institutional Animal Ethics Committee (Protocol numbers TIFRH/2019/02, TIFRH/2021/10). C57BL/6J and V2r1b-IRES-tauGFP mice, were purchased from JAX, housed and bred in individually ventilated cages, in a specific pathogen-free barrier facility, with a 12-hour light-dark cycle, ad-libitum provision of feed and water. Tunicamycin treatment was performed by intraperitoneal injection of 9-week-old C57BL/6J mice at a dose of 2 mg/kg body weight. Liver tissue was collected 48 hours after injection.

Cnpy-1 gene deletion mice were generated using CRISPR-Cas9 in the following manner. CRISPR gRNA sequence: ^5^’GATATTCGAACTTATTGCCC^3^’ was cloned into BbsI digested pX330 plasmid (Addgene# 42230). In vitro gRNA activity was assessed by transfecting N2A cells with the cloned gRNA, followed by T7E1 assay using standard methods (58). T7 sgRNA and Cas9 template were PCR amplified, gel purified, and in vitro transcribed, after which the purified RNA was microinjected into B6/CBA zygotes at a concentration of 50 ng/ul Cas9 and 25 ng/ul gRNA. One-cell fertilized embryos were injected into the pronucleus and cytoplasm of each zygote. Microinjections and mouse transgenesis experiments were performed as described previously (59, 60). Global knockout alleles were detected by PCR across the cleavage site, followed by T7E1 assay. Oligo sequences and band sizes: ∼280bp band cleaves to ∼180bp and ∼100bp.

Animals identified to contain the gene deletion allele by T7E1 assay were bred with C57BL/6J and offspring were analyzed by PCR, T7E1 and confirmed by Sanger sequencing. One of the lines resulting in a 99 base pair deletion (Chromosome 5: 28412242-28412340 as per mouse genome reference GRCm39) was confirmed for lack of Cnpy1 protein by anti-Cnpy1 labeling on VNO sections, western blot from VNO lysates. Mice containing this gene deletion allele were backcrossed for 8 generations to C57Bl/6J to generate a 98% pure background.

#### Antibody generation

Anti-peptide antibodies were generated by identifying amino acid sequence specific to the target protein and based on its predicted immunogenicity. KLH conjugated synthetic peptides were injected into rabbits to generate polyclonal antibodies, by YenZym Antibodies (Brisbane, CA). Antibodies were peptide affinity purified and dialyzed against PBS. Antibody specificity was confirmed via detection of target protein on tissue sections, as well as by blocking reactivity by pre-incubation with the peptide used for immunization (**Figure S14**). The following mouse peptide sequences from target proteins were used:

1. Cnpy1: DDPVTKQKYFRRYAPRKGD
2. Vmn2r111/Vmn2r112: FWKMKRNENKDRNQ
3. Vmn2r65: QKVNFTQKFSDTHSKIEYNH

Tissue Panel RT-PCR:

RNA was isolated from various tissues harvested from 12-weeks old C57BL/6J mice using the TRIzol reagent (Invitrogen #15596026) as per the manufacturer guidelines. cDNA was synthesized with 0.6ug of RNA from tissue using Superscript IV c-DNA synthesis kit (Thermo Fisher Scientific) with oligodT as per manufacturer instructions. cDNA was purified using a DNA cleanup kit (Monarch, NEB) and used as a template for performing PCR with primers targeting exon-1 or exon-2 of full-length *Cnpy1* and *Gapdh* as a positive control.

#### RNA sequencing and analysis

The full-length Cnpy1 transcript sequence was obtained using VNO RNA-seq FASTQ files from (19). to assemble the transcriptome. Raw reads were aligned to the mm10 reference genome using STAR (61), and the resulting BAM files were used as input for transcriptome assembly with StringTie (62).

VNO from Cnpy1^+/+^ and Cnpy1^-/-^ littermate animals (three each) were dissected. Lysis was performed in a Beadruptor using RLT buffer of Qiagen Rneasy plus kit with 1.4mm ceramic beads, and RNA was isolated as per manufacturer instructions. RNA was quantified using Qubit RNA BR assay kit and quality was checked using RNA Screen Tapes (Agilent, Cat#5067-5576)/RNA HS ScreenTapes (Agilent, Cat#5067-5579). 500ng RNA from each sample was used for library preparation using NEB Ultra II directional RNA-seq library prep kit (NEB #E7765S). Initially, poly-A containing mRNA were purified using oligo-dT magnetic beads from NEBNext Poly(A) magnetic isolation module (NEB #E7490). Following purification, the library was prepared as per manufacturer’s instructions. The libraries were indexed using NEBNext Multiplex Oligos for Illumina 96 Unique Dual Index Primer 96 rxn Set 3. Prepared libraries were quantified using Qubit HS Assay and pooled. Pooled libraries were sequenced on Illumina Novaseq instrument at 150bp paired end configuration to obtain 20Gb data per sample. Adaptor contamination was removed from the samples using trim_galore tool (https://www.bioinformatics.babraham.ac.uk/projects/trim_galore/) and Salmon tool (63) was used to quantify the gene expression. Raw counts from Salmon were used as input for DeSeq2 (64) to perform differential expression analysis between Cnpy1^-/-^ and Cnpy1^+/+^. pHeatmap R package was used to generate the heatmap of differentially expressed genes with an adjusted p-value < 0.01 and absolute fold change of 1.5.

#### qPCR

RNA isolation was done as described above. cDNA was prepared with 1ug of RNA using Superscript IV or Primescript cDNA synthesis kit and the same was used for qPCR reactions as a template. TaqMan probes (for V2Rs, *Gnao1* and Ddit3/CHOP genes) or gene specific primers (*Xbp1* spliced, unspliced) were used to set up reactions in triplicates using cDNA samples as templates as per the TaqMan master mix manual or FastStart essential DNA green mix. qPCR was performed on Roche lightCycler96 instrument, and data was exported, analyzed using the supplied software. Expression of each gene was normalized to *Gapdh* expression in that sample and represented as fold change relative to Cnpy1^+/+^ or Liver-control.

#### RNA *in situ* hybridization (ISH)

Chromogenic RNA ISH was performed as described before (15). Briefly, using VNO cDNA as template and primers containing T7 RNA polymerase promoter sequence, specific region of Cnpy1 coding sequence was PCR amplified. *In-vitro* transcription was performed using Digoxigenin (DIG) labeled UTP nucleotide mix and post RNA cleanup, riboprobes were stored at -80°C. Freshly dissected VNO was embedded in OCT and cryosectioned at 14µm thickness. Tissue sections captured on superfrost-plus slides were hybridized with Cnpy1 specific riboprobe, washed and hybridized probe was detected using anti-DIG conjugated with alkaline phosphatase. Colorimetric (BCIP/NBT) substrates were used to develop the bound antibody-enzyme conjugate reactions.

#### Cnpy1 expression in heterologous cells

Mouse Cnpy1 corresponding to the full-length sequence (Figure 1) was cloned by PCR amplification from VNO cDNA into pcDNA3.1+ or pcDNA4TO plasmids. Myc tag was added after the start codon or before the stop codon, to generate myc-Cnpy1 and Cnpy1-myc clones in the same plasmid vector. HEK293 cells, cultured under 5% CO_2_ in Dulbeco’s Modified Eagle Medium (DMEM) supplemented with GlutaMAX, penicillin-streptomycin and 5% FBS, were adhered to poly-D lysine coated 35mm glass coverslip bottom dishes. Plasmid DNA was transfected using Polyethylenimine transfection reagent. 18-24 hours post transfection; cells were fixed with 2% paraformaldehyde and expressed protein was detected using immunofluorescence microscopy.

#### Immunofluorescence microscopy

VNO tissues, either with or without outer bone, were harvested from animals, fixed for 10-20 minutes with 4% Paraformaldehyde in phosphate buffered saline (PBS). After washing off PFA, tissues with bone were treated with 0.5M EDTA for 2-16 hours (depending on mouse age) to soften the bone, followed by cryoprotection with 30% sucrose and embedding, freezing in OCT (Sakura) medium. 14µm sections were collected on superfrost-plus slides using a cryostat and fixed with 4% PFA. Unreacted PFA was quenched with 30mM glycine in PBS for 10 minutes. Sections were blocked and permeabilized using 3% bovine serum albumin (BSA) in 0.1% Triton-X 100 (0.1% Nonidet P40 for HEK293) detergent and labeled with primary antibodies diluted in the same blocking buffer. The respective primary antibodies were detected using Alexa Fluor 488, 647 or Cy3 fluorophore conjugated secondary antibodies. Images were acquired on Olympus FV3000, Leica Stellaris confocal microscopes using 10X-0.2NA air or 60X-1.4NA oil immersion objectives with sequential channel image acquisition. Images for quantification of Gαi2, Gαo zones, CC3, V2Rs or Ki67 fluorescence were acquired on Olympus IX83 inverted fluorescence microscope operating in widefield mode.

#### Image quantification

Fluorescent labeled cells from images of VNO sections (anti-CC3, anti-Ki67, anti-Vmn2r, anti-SEKDEL) were quantified automatically using ilastik tool (65). To avoid estimation errors caused by fluorescence from non-sensory epithelium, sensory epithelium was manually cropped in each section. Pixel classification in ilastik was trained on 10-15 representative images, using features including intensity and edge detection to differentiate truly labeled pixels from background. Next, object identification in ilastik was trained with pixel classification data to identify VSN cell bodies and exclude objects below a minimum size threshold. The program was also trained to classify single cells, doublets, or multiplets. Once identification was optimized separately for each antibody, batch processing was performed to produce data tables for each image. These tables were parsed with custom Python code, which summed singlets, doublets, and multiplets per image to calculate total cell counts. To quantify CC3 and SEKDEL double-positive cells, segmented masks of the SEKDEL-labeled area were generated using pixel classification workflow, applied to the CC3 channel, and the above quantification workflow was used for CC3+ cells. The area of SEKDEL labeling was measured from the segmented mask. Representative images for the steps in this process are shown in **Figure S15**. Quantification of Gαo, Gαi2 (anti-Nsg1) area in Figure 4 and pS6+ cells in Figure 8 was done manually.

#### Western Blot

VNOs pooled from 2-5 mice were lysed by passing through 10-20 strokes of a Dounce homogenizer using lysis (50 mM Tris - pH7.5, 150 mM NaCl, 1 mM EDTA, 1% Nonidet-P40, 1 mM PMSF, protease inhibitor cocktail, phosphatase inhibitor cocktail 2 and 3) on ice. To ensure complete homogenization, tissues were disrupted in Beadruptor, using 2.8 mm and 1.4 mm ceramic beads. Lysate was clarified by removing debris by centrifugation at 20,000xg at 4°C and used for total protein estimation via the BCA method (Thermo). Tissue lysate was mixed with Laemmli sample buffer, containing fresh DTT and heated at 90°C for 3 minutes before loading equal protein amount on a discontinuous 12% SDS-polyacrylamide gel. After gel run, proteins were transferred to 0.2 μm PVDF membrane in Tris-Glycine transfer buffer with 10% methanol. Membranes were blocked with 5% non-fat milk or Starting Block (Thermo) in Tris buffered saline containing 0.1% Tween-20. Blocked membranes were incubated with primary antibodies and HRP conjugated secondary antibodies. Chemiluminescence was detected using ECL Plus kit (Cytiva) on a chemidoc (Vilbur instruments). Quantification of signal was performed by “Gels” analysis module from ImageJ/Fiji (66). Quantification was performed using non-saturating exposures and target protein levels were normalized to β-tubulin as internal loading control.

#### VNO lysate and immunoprecipitation (IP)

IP_Cnpy1:B6_ and IP_Cnpy1:KO_, were performed as three technical replaces, and IP_IgG:B6_ as 2 replicates. Each replicate comprised of VNOs collected from 50 mice of 8-16 weeks age and roughly equal numbers of male and female. VNOs were dissected, pooled and kept frozen at -80°C. On the day of experiment, tubes containing VNOs were thawed and transferred to a 2ml Dounce homogenizer with cold lysis buffer. Homogenization was performed as described above. Cell debris was pelleted by centrifugation at 15000 rpm at 4°C for 15 min and supernatant was transferred to a new tube. After 2 hours of preclearing with Protein-A Mag Sepharose beads, 15µg of Cnpy1 antibody or normal rabbit IgG was added to each tube. After 16 hours of antibody incubation on end-to-end rotator at 4°C, antibodies were precipitated by adding mag-sepharose beads for 4 hours. Magnetic beads containing antibody-protein complex were retained using magnetic racks, washed three times with lysis buffer. Elution of antibodies from Protein-A beads was performed with 2X Laemmli buffer at 95°C. The eluate was used for mass spectrometry.

#### Sample preparation for quantitative mass spectrometry (MS)

The SP3 protocol (67, 68) was performed on a KingFisher system (Thermo) for sample clean-up. Trypsin (sequencing grade, Promega) was added in an enzyme to protein ratio 1:20 for a 5h digestion at 37°C (in 50 mM HEPES) together with 5mM tris(2-carboxyethyl)phosphine and 20mM 2-chloroacetamide.

Peptides were labelled with TMT10plex isobaric labeling reagent (Thermo) (69) according to the manufacturer’s instructions. In brief, of 0.8mg reagent dissolved in 42ul acetonitrile (100%) 8 ul was added and incubated for 1h room temperature. The reaction was stopped with 8 ul 5% hydroxylamine for 15 minutes at RT. Samples of a set were combined and desalted on a OASIS® HLB µElution Plate (Waters). Offline high pH reverse phase fractionation was carried out on an Agilent 1200 Infinity high-performance liquid chromatography system, equipped with a Gemini C18 column (3 μm, 110 Å, 100 x 1.0 mm, Phenomenex) installed with a Gemini C18, 4 x 2.0 mm SecurityGuard (Phenomenex) cartridge as a guard column. The binary solvent system consisted of 20 mM ammonium formate (pH 10.0) (A) and 100% acetonitrile as mobile phase (B). The flow rate was set to 0.1 mL/min. Peptides were separated using a gradient of 100% A for 2 min, to 35% B in 59 min, to 85% B in another 1 min and kept at 85% B for an additional 15 min, before returning to 100% A and re-equilibration for 13 min. 48 fractions were collected which were pooled into 6 fractions. Pooled fractions were dried under vacuum centrifugation, reconstituted in 10 μL 1% formic acid, 4% acetonitrile and then stored at -80 °C until LC-MS analysis.

#### MS data acquisition

An UltiMate 3000 RSLC nano LC system (Dionex) fitted with a trapping cartridge (µ-Precolumn C18 PepMap 100, 5µm, 300 µm i.d. x 5 mm, 100 Å) and an analytical column (nanoEase™ M/Z HSS T3 column 75 µm x 250 mm C18, 1.8 µm, 100 Å, Waters) was coupled to an Q-Exactive Plus Mass Spectrometer (Thermo) using the Nanospray Flex™ ion source in positive ion mode.

The samples were loaded onto the trapping column with a constant flow of 30 µL/min 0.05% trifluoroacetic acid in water for 4 minutes. After switching in line with the analytical column peptides were eluted at a constant flow of 0.3 µL/min using the method described in the following. The binary solvent system consisted of 0.1% formic acid in water with 3% DMSO (solvent A) and 0.1% formic acid in acetonitrile with 3% DMSO (solvent B). The percentage of solvent B was increased from 2% to 8% in 2 min, from 8% to 28% in 66 min, from 28%-40% in another 10 min and finally from 40%-80% in 2.5 min, followed by re-equilibration back to 2% B in 5.5 min.

The peptides were introduced into the MS instrument via a Pico-Tip Emitter 360 µm OD x 20 µm ID; 10 µm tip (CoAnn Technologies) and an applied spray voltage of 2.4 kV. The capillary temperature was set at 275°C. Full mass scan was acquired with a mass range 375-1200 m/z in profile mode in the orbitrap with resolution of 70000. The fill time was set at a maximum of 250 ms with a limitation of 3x10^6^ ions. Data dependent acquisition (DDA) was performed with the resolution of the Orbitrap set to 35000, with a fill time of 120 ms and a limitation of 2x10^5^ ions. A normalized collision energy of 32 was applied. MS^2^ data was acquired in profile mode. Define first mass was set to 110 m/z.

#### MS - data analysis

Fragpipe v21.1 with MS Fragger v4.0 (70) was used to process the acquired data, which was searched against a mus musculus proteome database (UP000000589, 21968entries, October 2022) plus common contaminants and reversed sequences. The following modifications were included into the search parameters: Carbamidomethyl on Cysteine and TMT10 on lysine as fixed modifications, protein N-term acetylation, oxidation on methionine and TMT10 on N-termini as variable modifications. A mass error tolerance of 20 ppm was used for precursor as well as for fragment ions. Trypsin was set as protease with a maximum of two missed cleavages. The minimum peptide length was set to seven amino acids. The false discovery rate on peptide and protein level was set to 0.01.

The raw output files of FragPipe (protein.tsv files) were processed using the R programming language. Contaminants and reverse proteins were filtered out and only proteins that were quantified with at least 2 razor peptides (Razor.Peptides >= 2) were considered for the analysis. 1980 proteins passed the quality control filters. Log2 transformed raw TMT reporter ion intensities (’channel’ columns) were first cleaned for batch effects using the ’removeBatchEffect’ function of the limma package (71) and further normalized using the ’normalizeVSN’ function of the limma package (72). Proteins were tested for differential expression using a moderated t-test by applying the limma package (’lmFit’ and ’eBayes’ functions). The replicate information was added as a factor in the design matrix given as an argument to the ’lmFit’ function of limma. A protein was annotated as an enriched hit with a false discovery rate (fdr) smaller than 0.05 and absolute fold change >= 2; or as a hit with a fdr below 0.05 and an absolute fold change between 1.5 and 2.0 in each comparison. The common enriched hits between IP_Cnpy1:B6_ vs IP_IgG:B6_ and IP_Cnpy1:B6_ vs IP_Cnpy1:KO_ were considered as potential interacting proteins of Cnpy1.

#### Rat and opposite sex bedding exposure

Male Cnpy1^+/+^ and Cnpy1^-/-^ mice aged 10–11 weeks were individually housed and isolated for a minimum of three days. Following isolation, soiled bedding collected from three male Wistar rats housed together was introduced into the home cage of each mouse. After one hour of exposure to rat bedding, the mice were euthanized, and VNOs were dissected. Female or male bedding was freshly collected from group housed adult females and the same procedure was followed. The VNO tissues were fixed in 4% paraformaldehyde for four hours and subsequently cryoprotected with 30% sucrose overnight. Immunohistochemistry for pS6 and SEKDEL was performed according to the procedure described above. Cells positive for pS6 within SEKDEL-labeled regions or throughout the neuroepithelium were counted manually.

#### Resident-intruder assay

Castration of C57Bl/6J mice was performed by orchidectomy procedure, performed on 6–8 weeks old C57BL/6J males chosen to be 10% lighter in weight than residents. Mice were anesthetized with Ketamine-Xylazine, followed by Meloxicam and Ceftriaxone-Tazobactam. After making a midline incision in the scrotum, both testes were exteriorized, ligated and removed. The incision was closed with sutures, iodine and antiseptic powder were applied, and mice were allowed 10 days for recovery.

Cnpy1^+/+^, Cnpy1^+/-^ and Cnpy1^-/-^ male mice from Cnpy1^+/-^ x Cnpy1^+/-^ of 10-12 weeks age were individually co-housed with a female for five days, followed by removal of the females. Starting one hour after the onset of the dark phase, castrated C57BL/6J males (intruder), swabbed with fresh urine from intact C57BL/6J males, were introduced into the home cage of the resident male for 10 minutes daily over four consecutive days, comprising the training phase. On the fifth day, resident-intruder interactions were recorded under infrared illumination using two Hikvision CCTV cameras from the top and side of the resident’s home cage. Aggressive behaviors, including attacks, chases, and wrestling, were manually scored from within the first 10 minutes of the video recordings. Resident and intruder males were separated after 15 minutes, or earlier, in case of extremely aggressive interactions. Scoring and quantification of aggressive behaviors was done by blinding the genotype. Two Cnpy1^+/+^ males that did not show any aggressive behavior during the training phase were excluded.

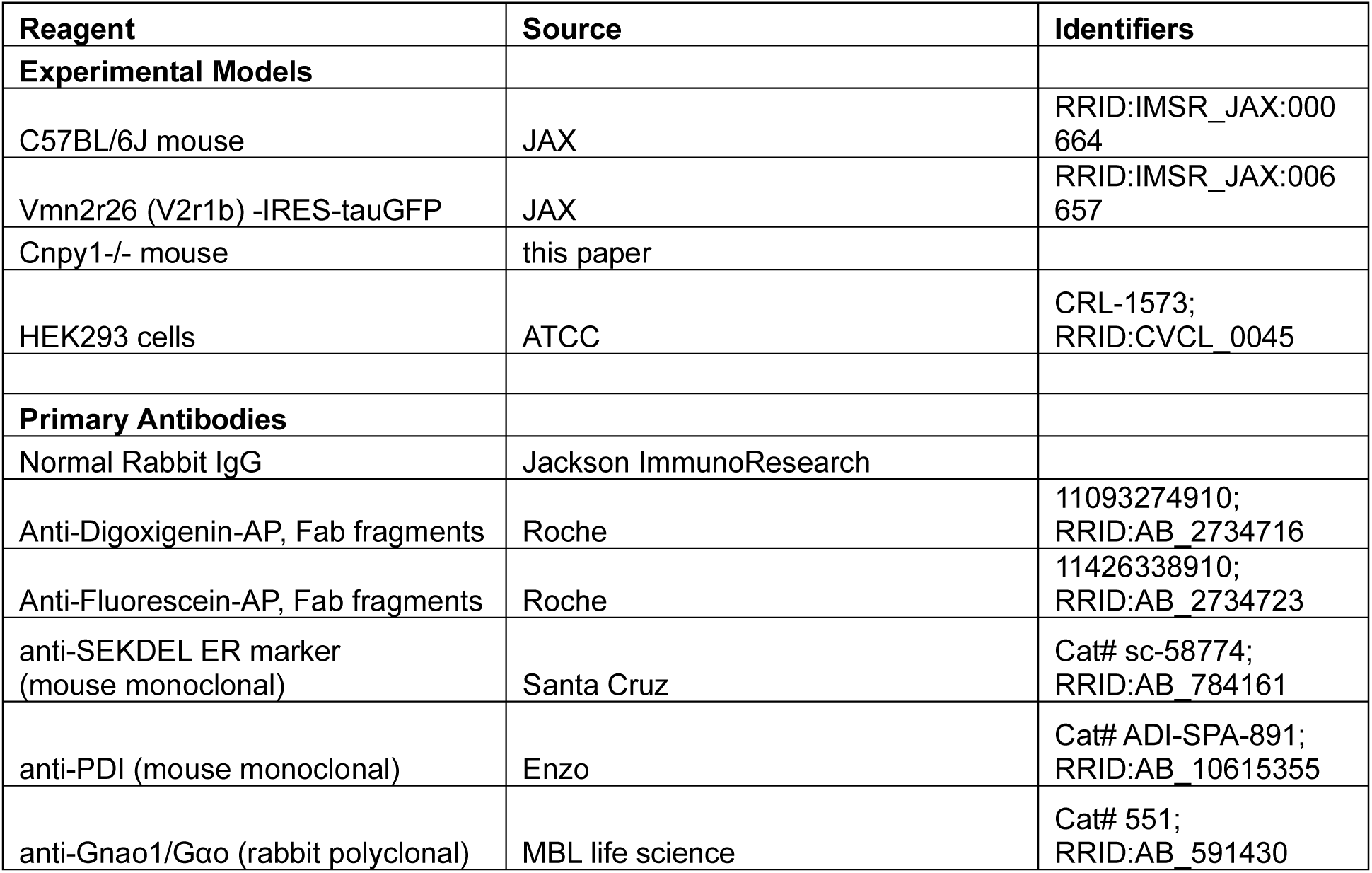
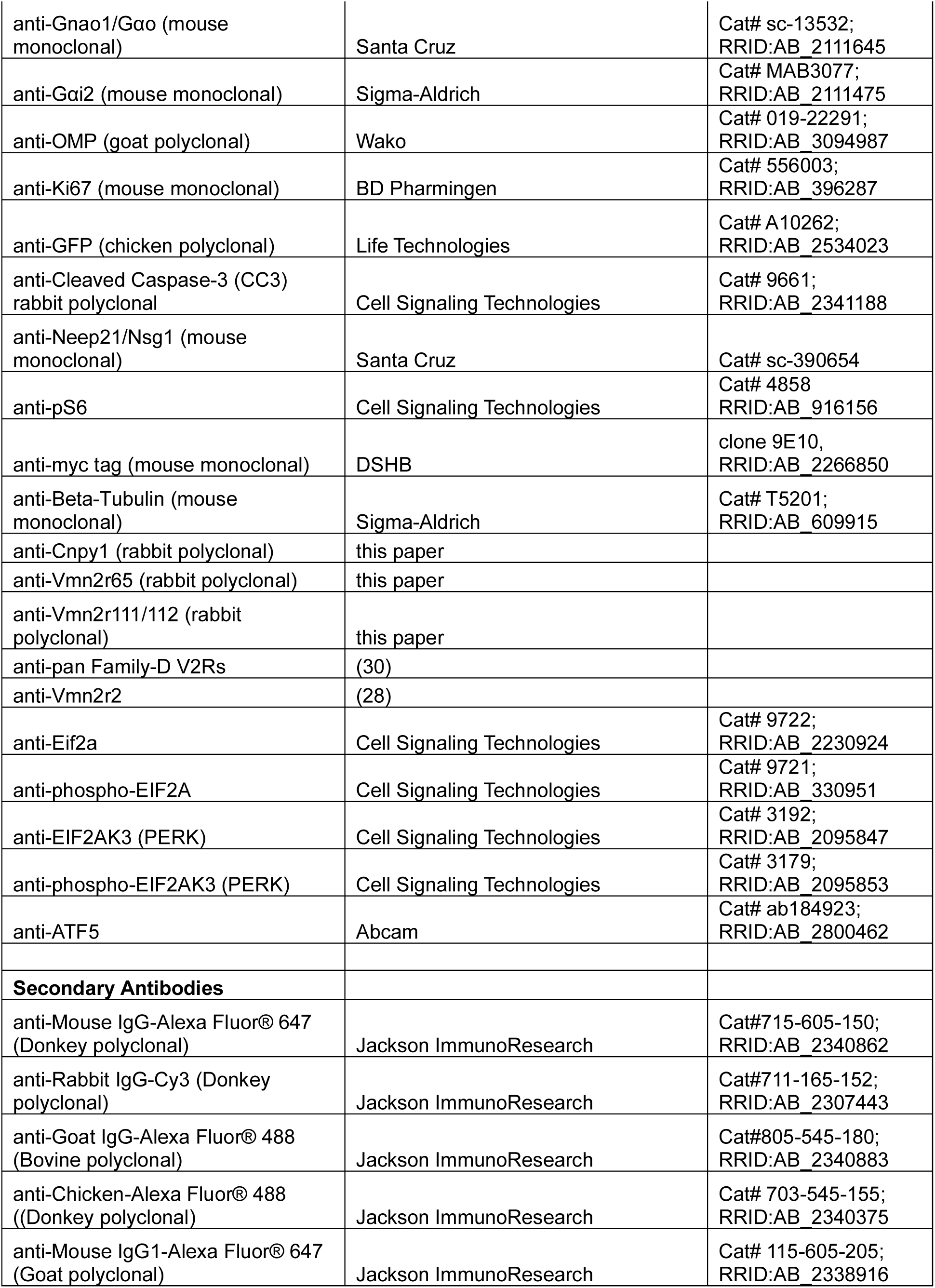
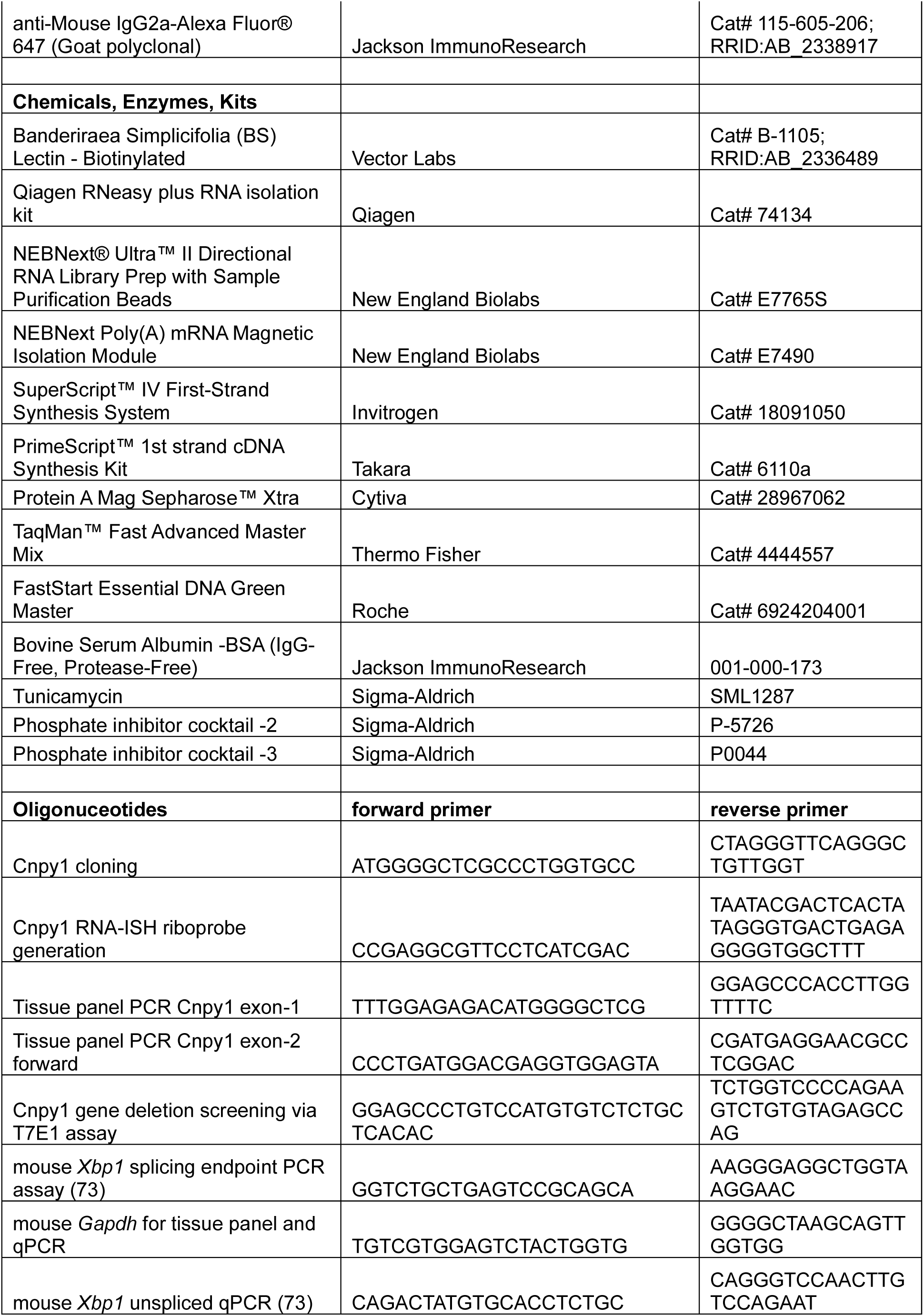
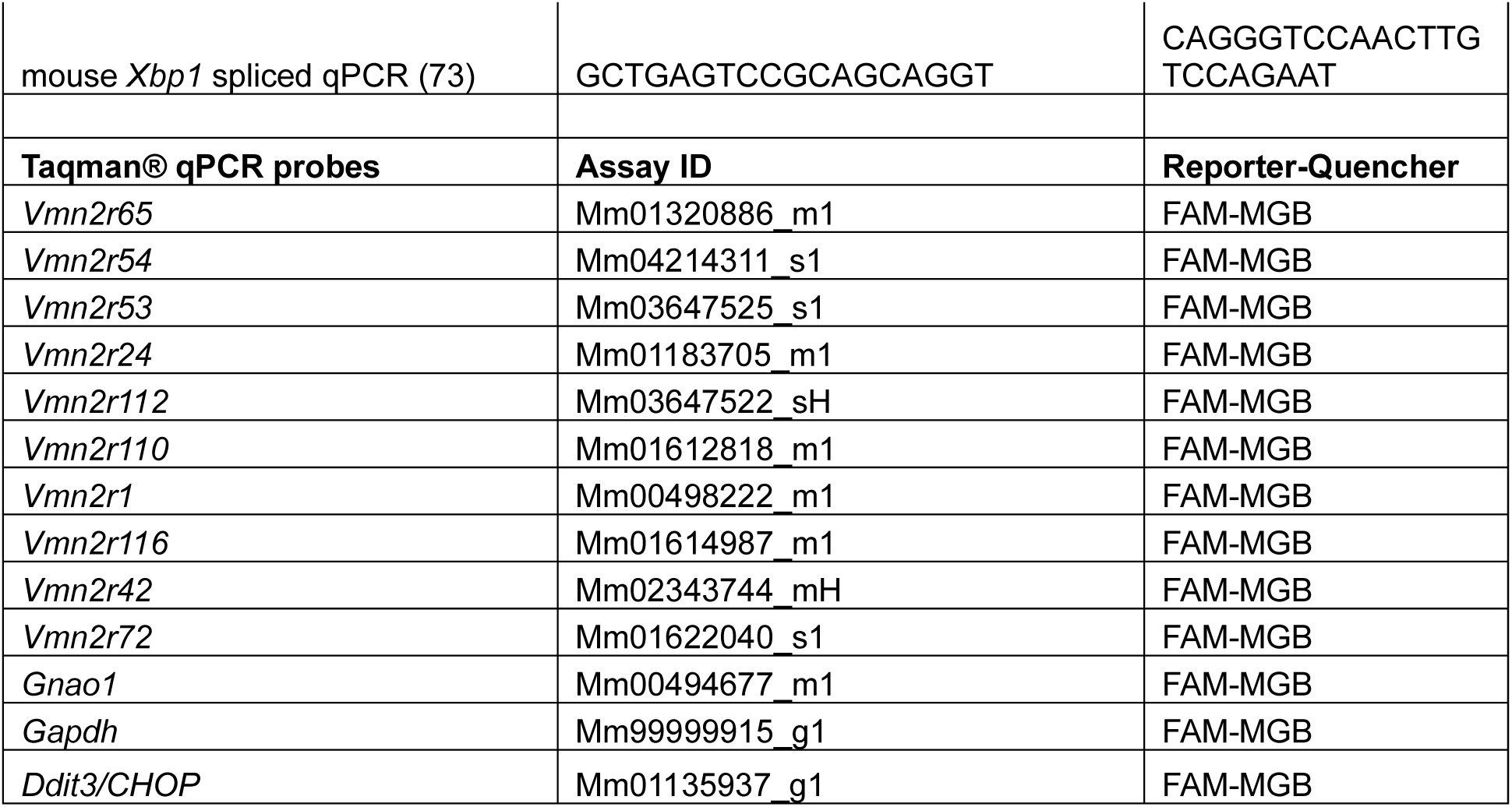

## Acknowledgements

We acknowledge part support from the Hope Center viral vectors and Mouse Genetics Core facilities at Washington University School of Medine, St. Louis, MO in generation of Cnpy1 gene deletion mouse model; We thank Jennifer Schwarz and Frank Stein of the Proteomics Core Facility at EMBL Heidelberg for help with peptide mass-spectrometry and analysis; Roberto Tirindelli for anti-pan family-D, anti-Vmn2r2 V2R antibody; Sritama Datta for technical assistance, Gopalkrishna R. for training with mouse surgery, Ganesh Kadasoor for CellSens software. This work was funded by Department of Atomic Energy, Government of India (Project No RTI 4007) and NIH Grant R21 MH099798.

## Data Availability

The following data are deposited to publicly accessible repositories: Mouse *Cnpy1* transcript sequence GenBank ID PX423936, RNA sequencing GEO ID GSE313188, IP-mass spectrometry proteomics: ProteomeXchange Consortium PRIDE repository ID: PXD068826.

---

**Figure S1.**
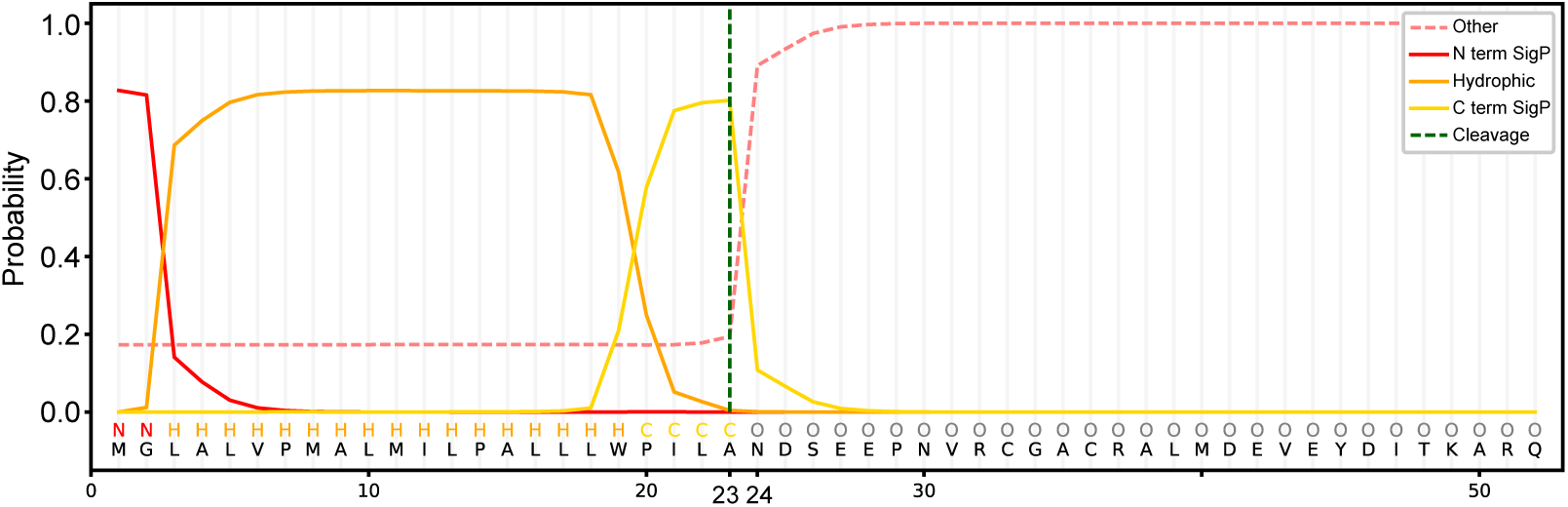
In silico identification of signal peptide sequence in full-length mouse Cnpy1. Cnpy1 N-terminal signal sequence predicted by SignalP 6.0 with cleavage site between amino acids 23-24 of mouse Cnpy1-FL indicated by vertical green dashed line. The probability plot shows signal peptide likelihood, including hydrophobic h-region preced-ing the cleavage site. Signal peptide cleavage is experimentally validated in **Figure S3**

**Figure S2.**
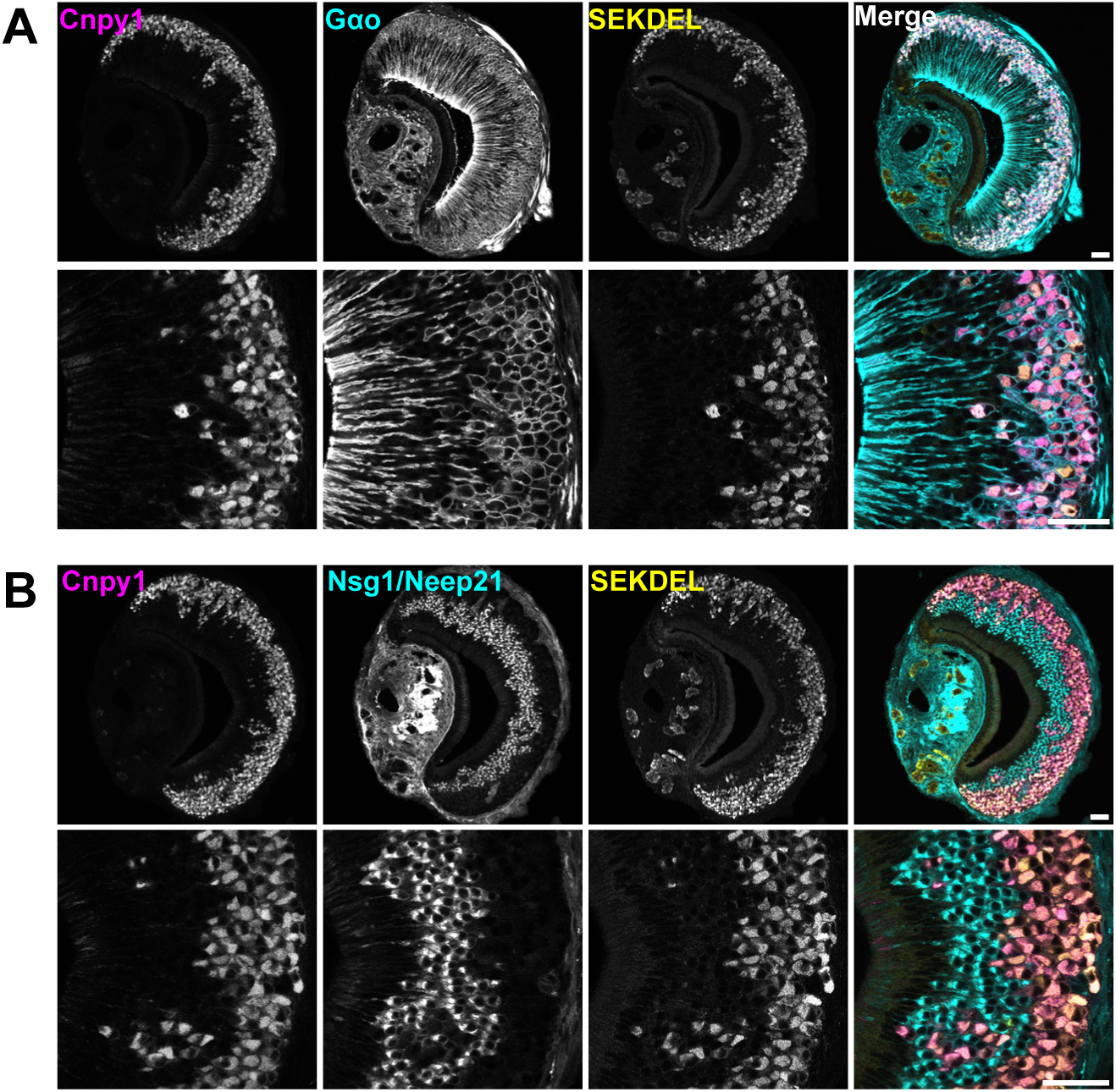
Cnpy1 is expressed in Gαo neurons. **A**) Immunofluorescence confocal microscopy images of VNO sections labelled with anti-Cnpy1, anti-Gαo, and anti-SEKDEL; show SEKDEL and Cnpy1 co-localization within Gαo neurons. Higher magnification images are shown in the bottom panel. **B**) Images of VNO sections labelled with anti-Cnpy1, anti-Nsg1/Neep21 (marker for Gαi2 neurons), anti-SEKDEL confirm the exclusive expression of Cnpy1 in Gαo neurons. Scale bar: 50μm.

**Figure S3.**
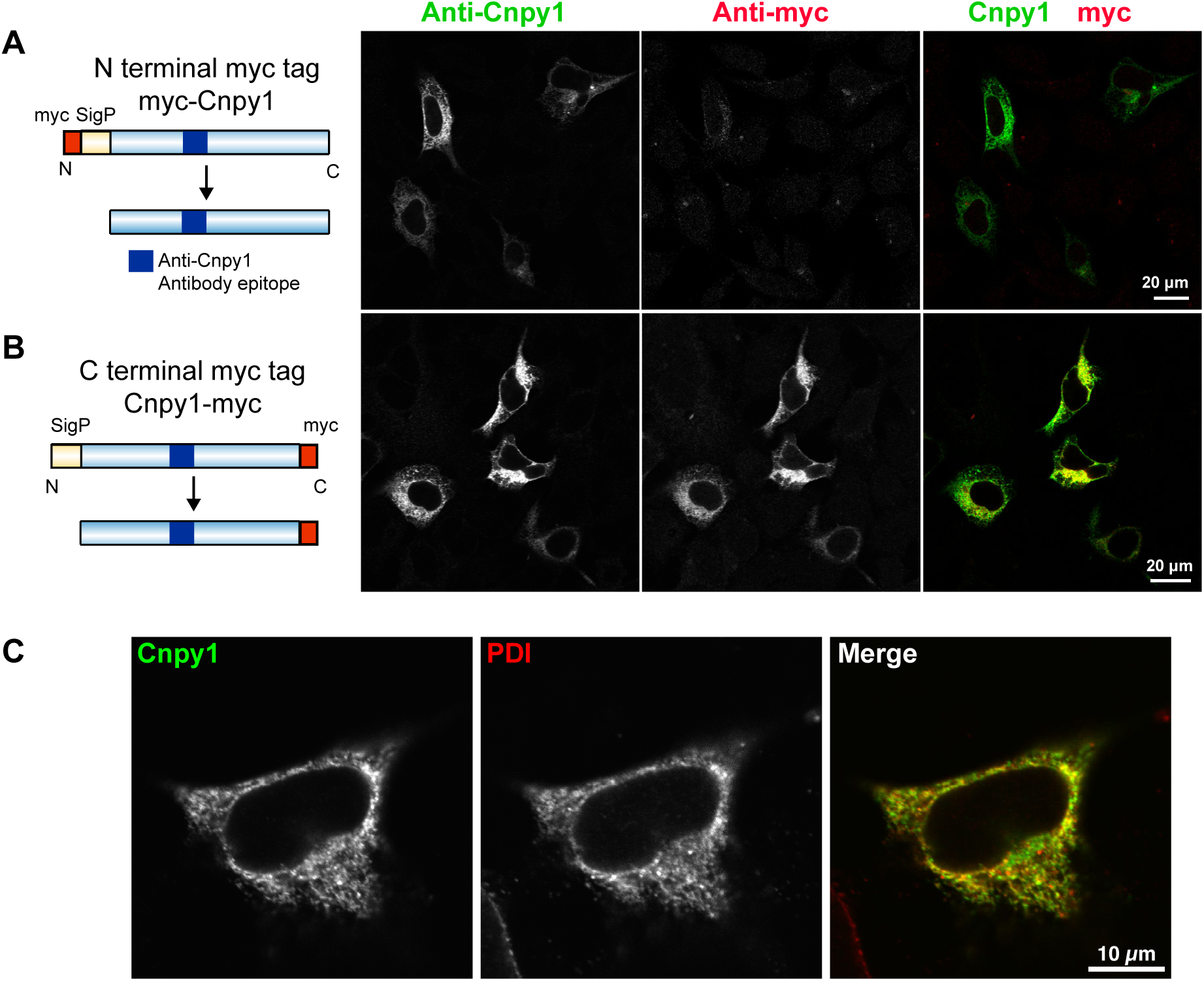
Expression of mouse Cnpy1 in HEK293 cells. **A, B**) Cnpy1-FL cloned with N-terminal myc tag (myc-Cnpy1) (**A**) and C-terminal myc tag (Cnpy1-myc) (**B**), were transiently expressed in HEK293 cells and immunolabeled with anti-Cnpy1, anti-myc tag. The schematic on the left shows the position of myc with respect to signal peptide (SigP) and location of Cnpy1 antibody binding epitope. Anti-Cnpy1 detects the expression of both constructs, whereas anti-myc signal is seen only with Cnpy1-myc, confirming the cleavage of N-terminal signal peptide. **C**) Immunolabeling of transiently expressed Cnpy1 and an ER resident protein – PDI shows ER localization in HEK293 cells.

**Figure S4.**
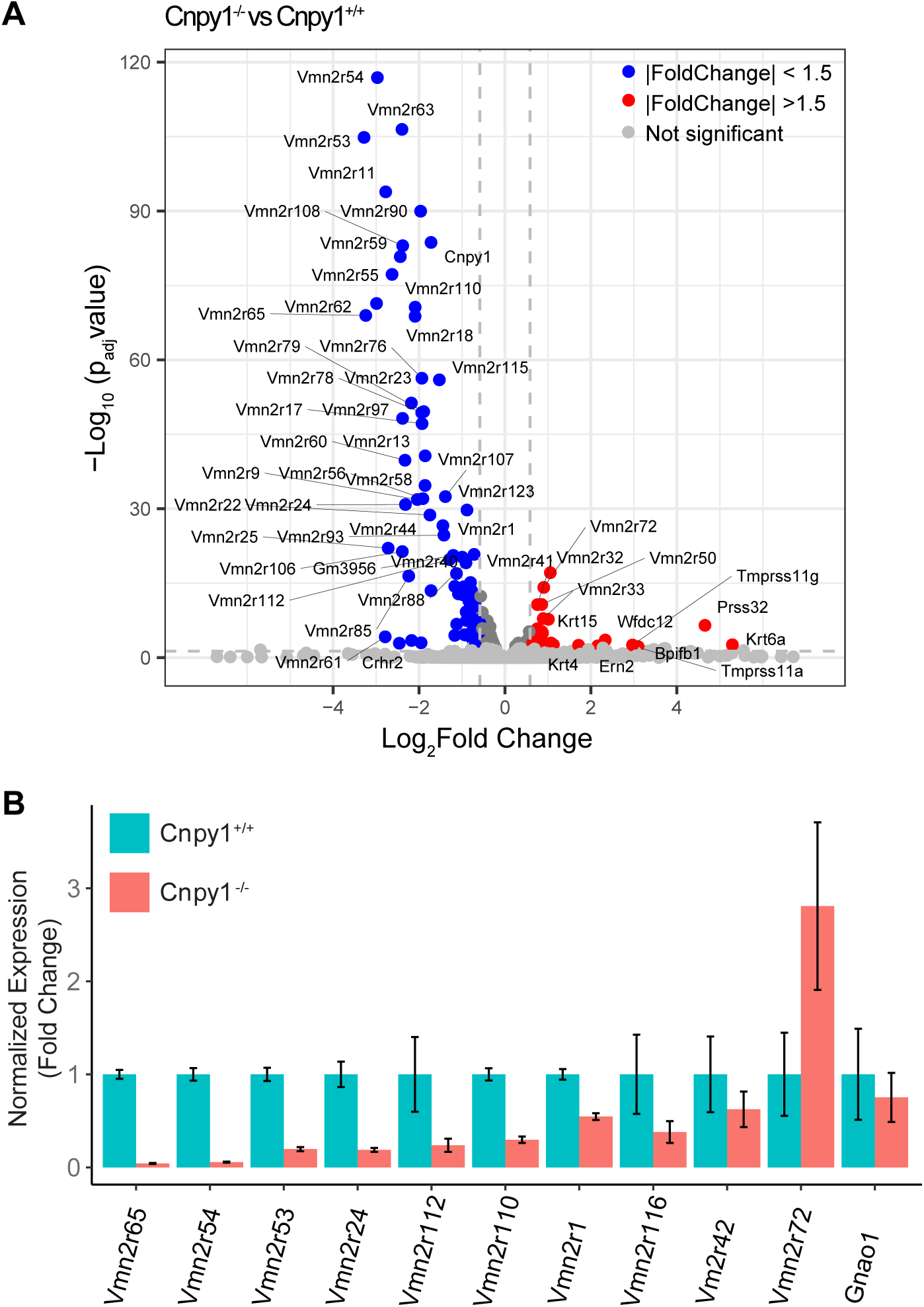
Gene expression differences between Cnpy1^+/+^ and Cnpy1^-/-^ VNO. **A)** Volcano plot (corresponding to Figure 3D), comparing gene expression (n =3 samples per genotype). The x-axis indicates the log2 fold change, and the y-axis represents the -log10 of the adjusted p-value (padj). Genes with significantly reduced transcripts in Cnpy1^-/-^ are shown in blue, and significantly upregulated ones are shown in red. Horizontal and vertical dashed lines indicate the significance thresholds (padj < 0.01 and |fold change| >1.5, respectively). Gray dots represent genes that did not meet the significance criteria. A complete list of differentially expressed genes is available in **Supplementary Table 1**. **B**) Validation of differences observed in Cnpy1^-/-^ and Cnpy1^+/+^ VNO bulk RNA sequencing using TaqMan probes specific for selected Vmn2r genes. Expression level of each gene relative to Cnpy1^+/+^ is represented as mean and error bars represent standard error.

**Figure S5.**
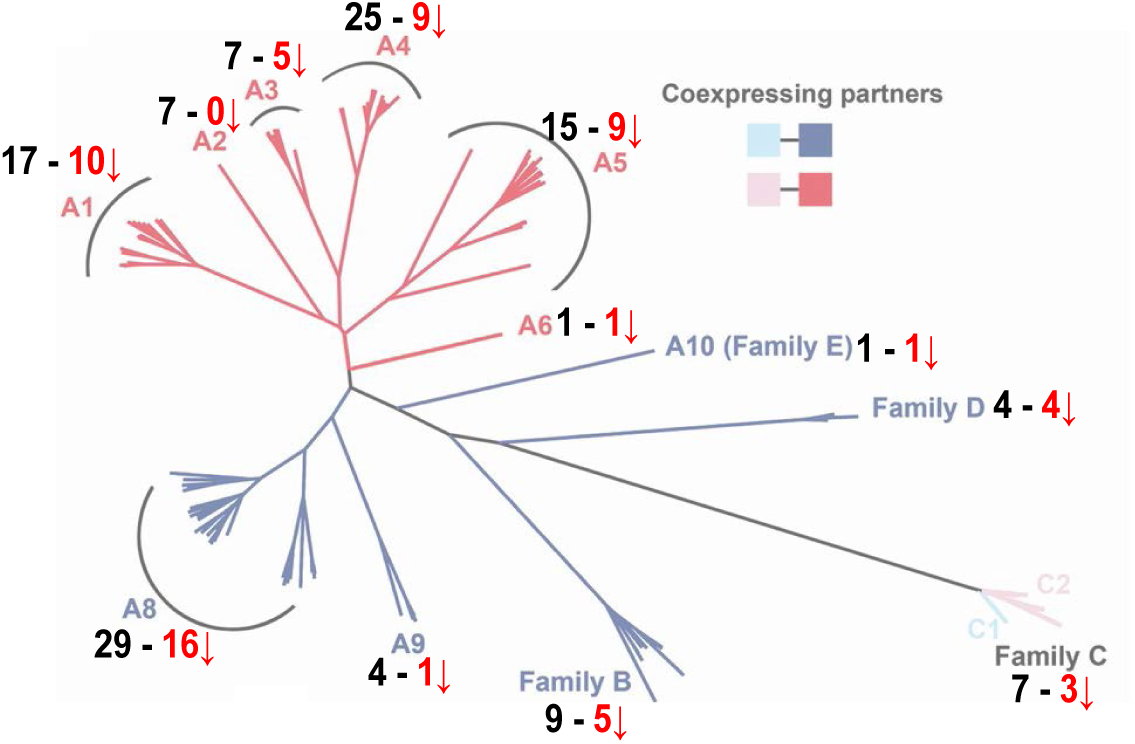
V2R downregulation in Cnpy1 knockout is not family-specific. Phylogenetic tree adapted from *Francia et al., 2014* (licensed under CC BY), showing classification of V2Rs into families based on sequence homology. The number of downregulated V2Rs identified in Figure 3D is shown in red with a downward facing arrow, adjacent to the total count for each family. Downregulated V2Rs are distributed across all families.

**Figure S6.**
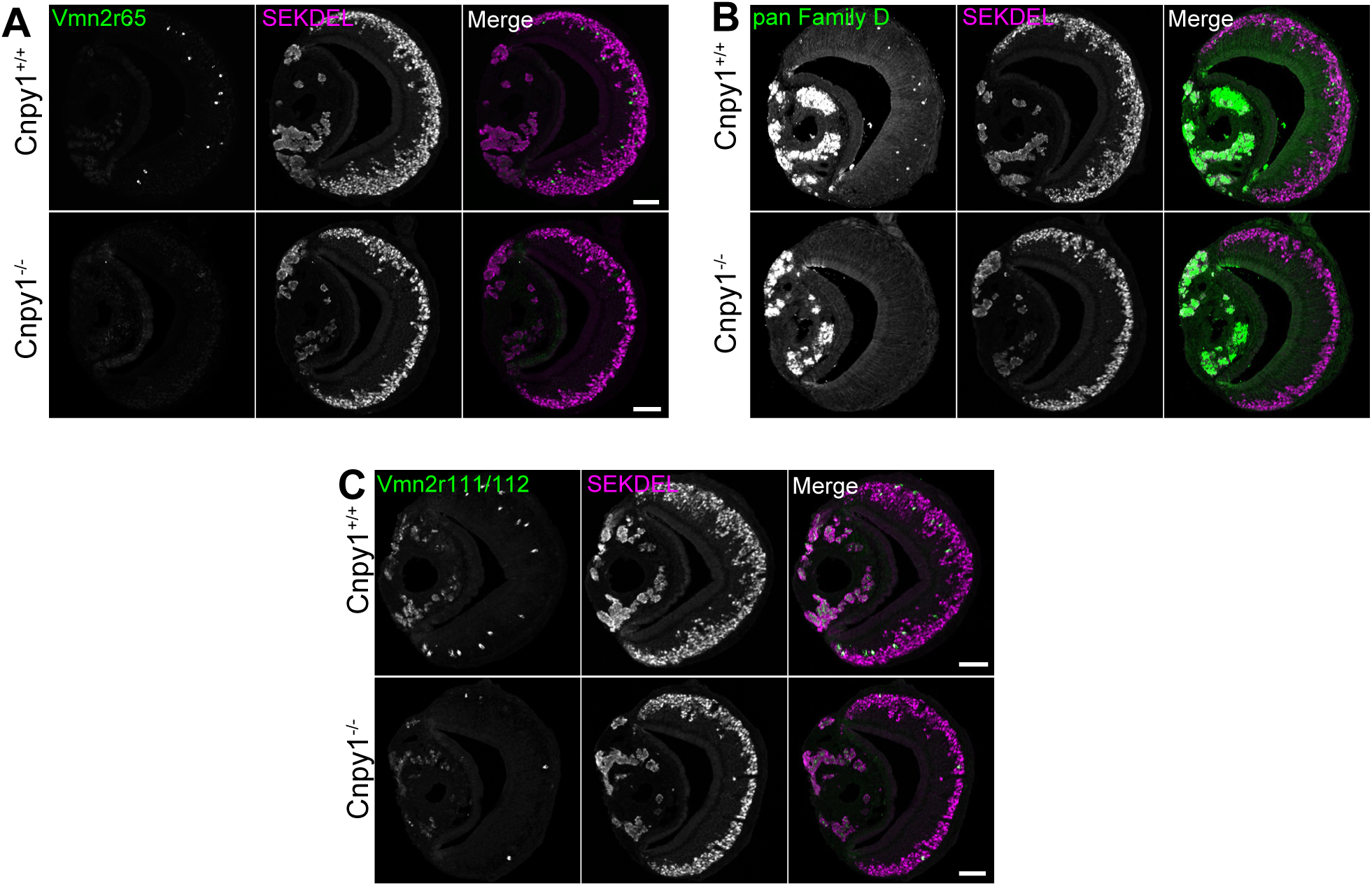
Immunolabeling of p60 VNO sections with antibodies against Vmn2r65 (**A**), pan-family D V2Rs (**B**), Vmn2r111/112 (**C**) shows the absence of these V2R expressing neurons in Cnpy1^-/-^ VNOs compared to Cnpy1^+/+^ as shown in **Figure 3**. Sections were co-labeled with an ER marker SEKDEL as a positive control to confirm tissue integrity and success-ful immuno-labeling. Scale bar: 100 μm

**Figure S7.**
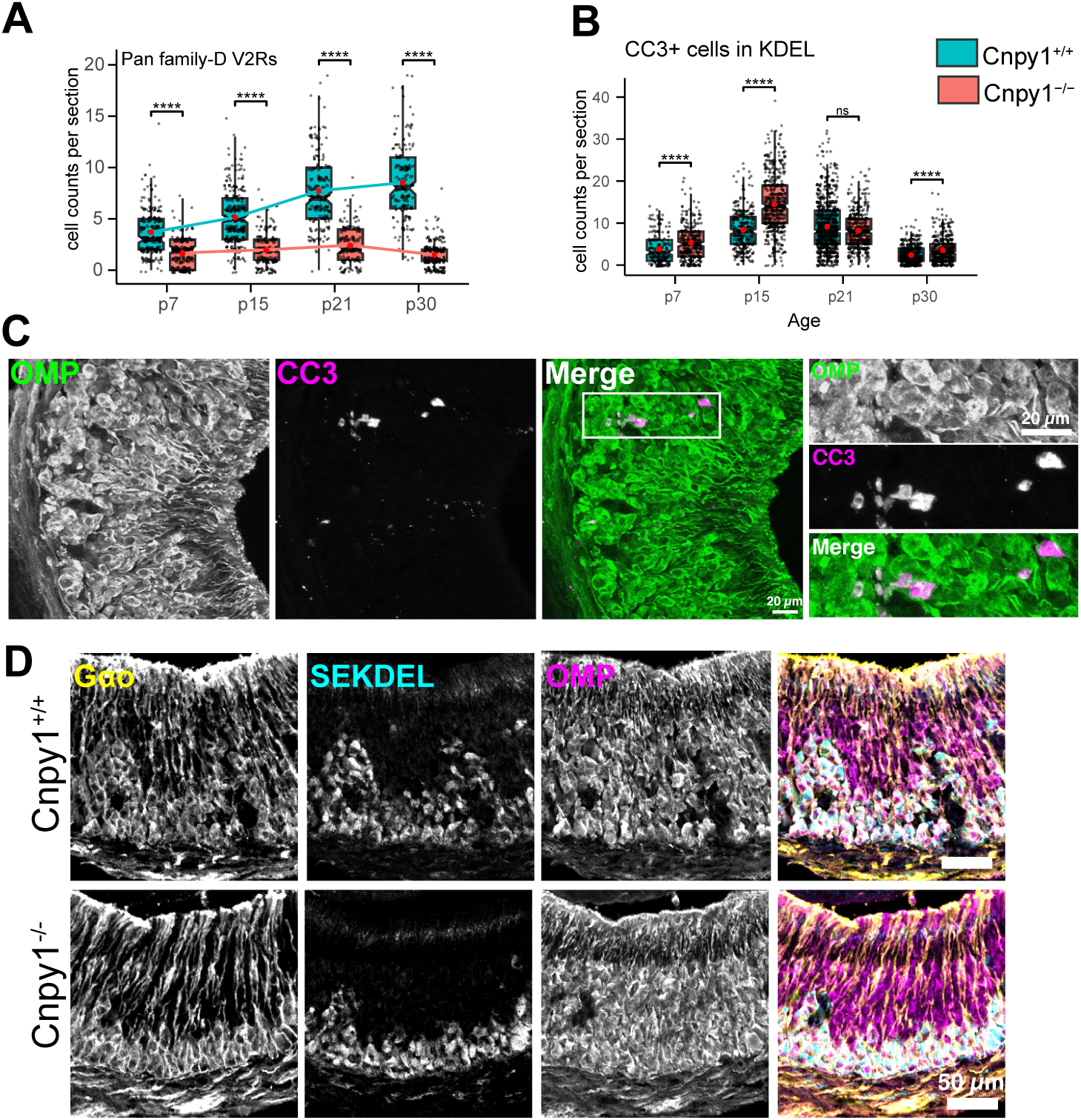
A) Cell counts per section from VNO sections across postnatal age labeled with anti-pan family-D V2Rs show progressive decline in Family-D expressing neurons during postnatal development of Cnpy1^-/-^ mice. **B**) Total number of CC3 labeled cells overlapping with Gαo (anti-SEKDEL) zone per section, quantified across postnatal age. Mann–Whitney U test was performed to measure statistical significance. **** - p value < 10-4 and ns - not significant. **C**) Immunolabeling of OMP and CC3 on p15 Cnpy1^-/-^ sections showing CC3+ apoptotic neurons are also labeled with OMP, a marker of mature neurons. **D**) Representative immuno-fluorescence confocal micros-copy images of VNO sections from p60 mice, labelled with anti-Gαo, anti-SEKDEL and anti-OMP. Similar to Cnpy1^+/+^ VNO, anti-SEKDEL is a marker for Cnpy1^-/-^ Gαo neurons. Gαo neurons in Cnpy1^-/-^ are also labelled with anti-OMP indicating their developmental status as mature neurons.

**Figure S8.**
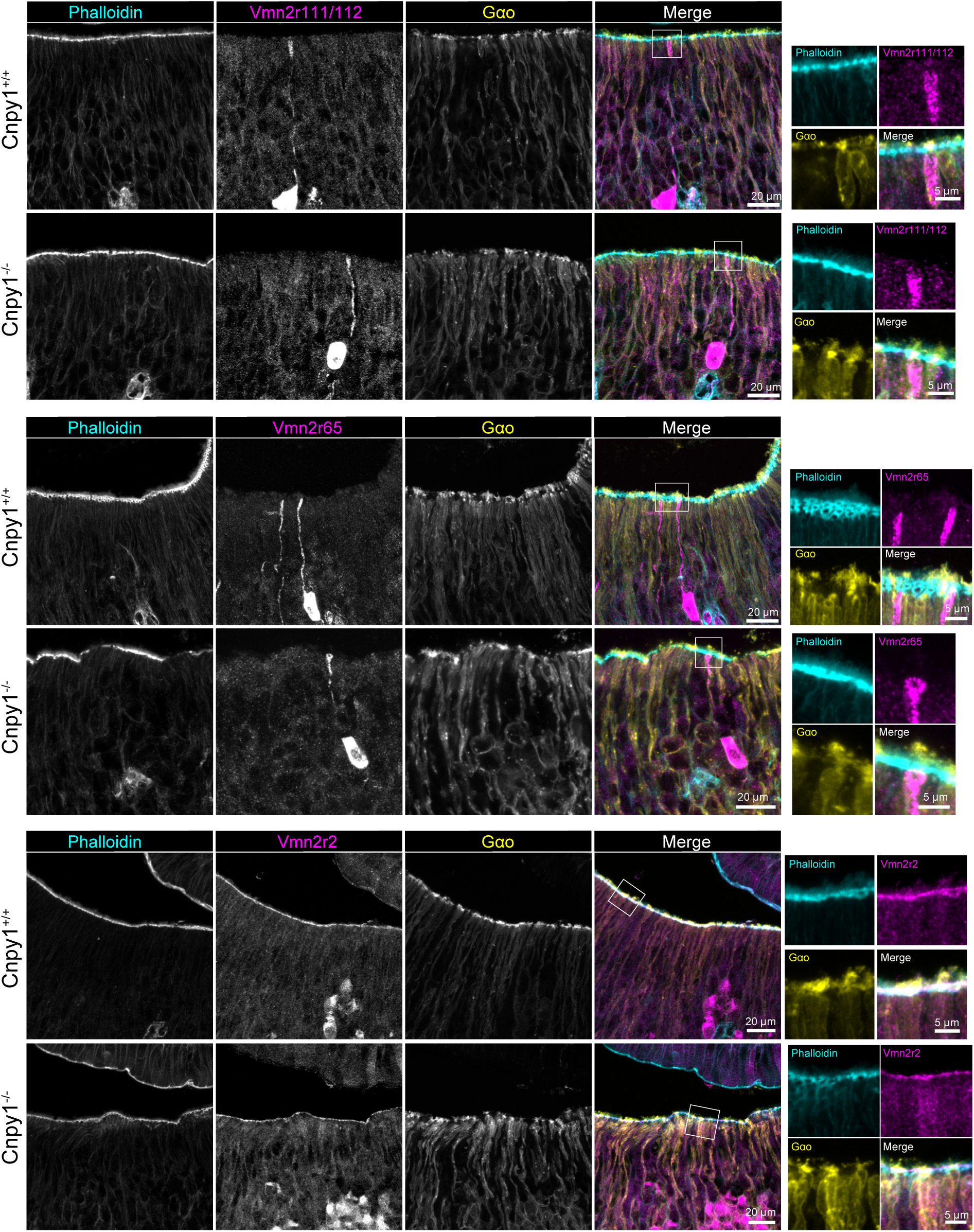
V2R trafficking and localization to dendritic tip are preserved in Cnpy1^-/-^ VSNs. VNO sensory epithelium from P14 Cnpy1^+/+^ (top) and Cnpy1^-/-^ (bottom) imaged towards the luminal interface by confocal microscopy. VNO sections are labelled for Phalloidin (cyan) - which marks VSN microvilli, anti-V2Rs (magenta) and anti-Gαo (yellow). In both genotypes, V2R signal is seen to track the projection and localize to the microvillar sensory interface, demonstrating that the transport of V2R proteins is not affected by the loss of Cnpy1.

**Figure S9.**
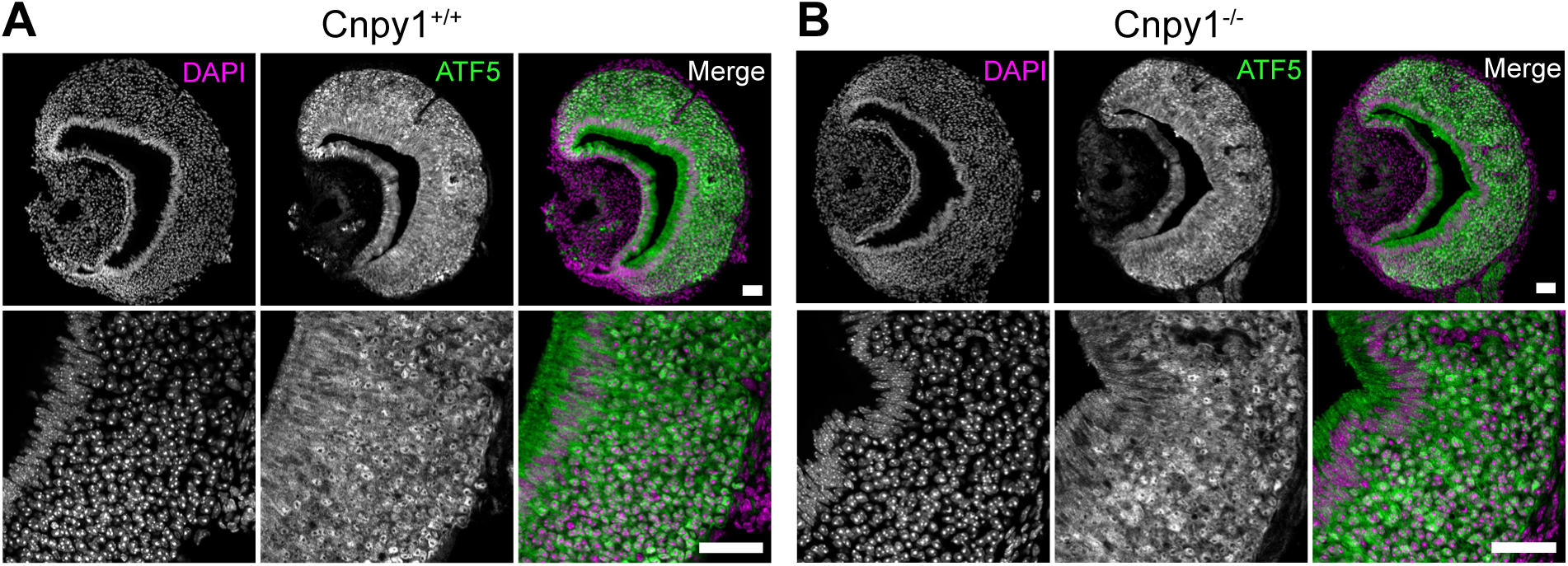
No change in VNO ATF5 expression in the absence of Cnpy1 **A-B**) Representative images from P14 Cnpy1^+/+^ (**A**) or Cnpy1^-/-^ (**B**) VNO sections immunolabeled for ATF5. Bottom panels show cellular detail of the sensory epithelium. ATF5 signal intensity or distribution is similar across both genotypes. Scale bar: 50 μm.

**Figure S10.**
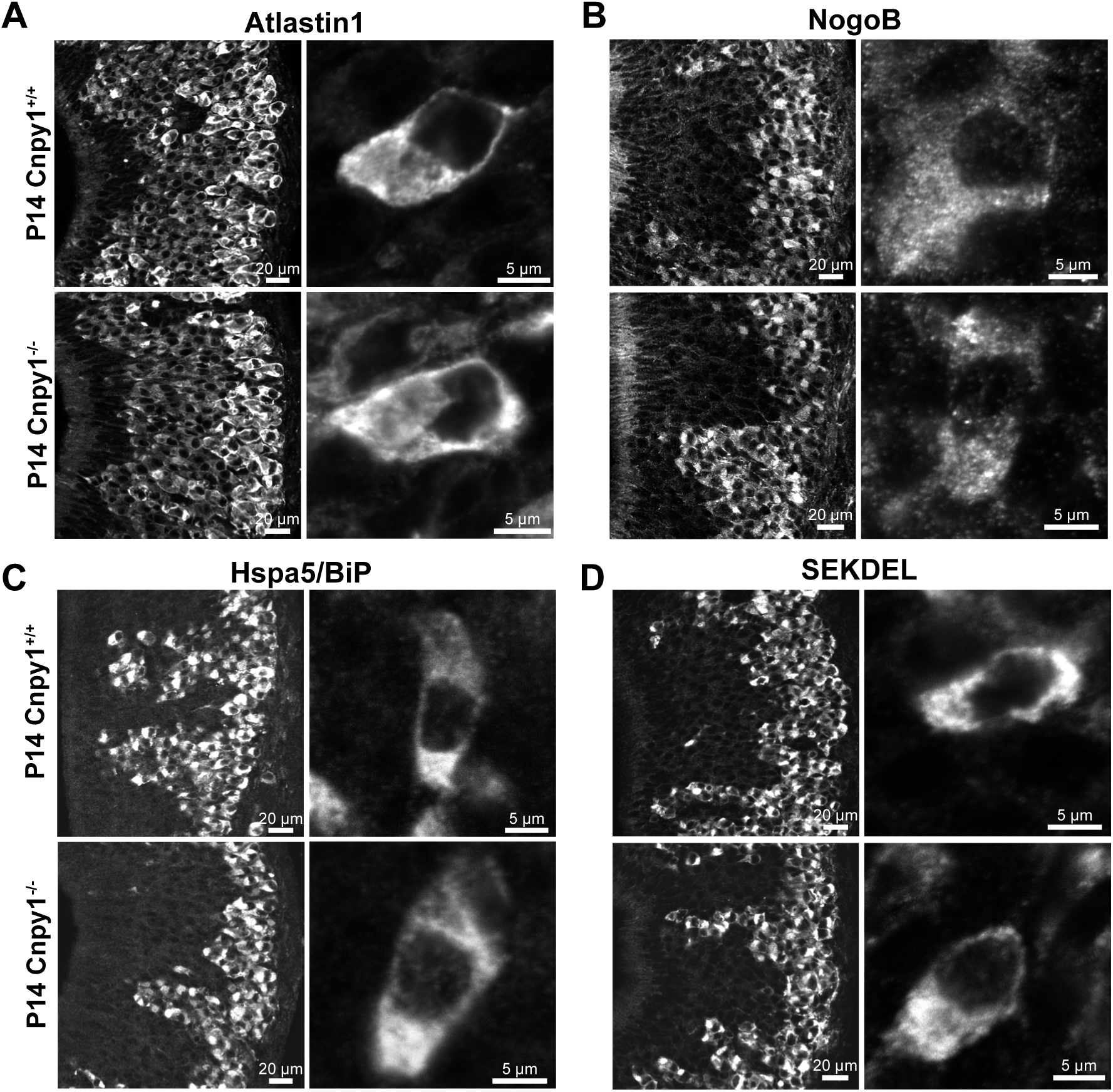
ER protein enrichment and ER structure is not affected by loss of Cnpy1. **A-D**) Confocal microscopy immuno-fluorescence images of VNO sections labelled with antibodies to ER structural proteins - Atlastin1 (**A**) and NogoB (**B**); the ER chaperone - Hspa5 (**C**); and ER marker SEKDEL **(D)**. For each panel, a low-magnification overview (left) shows the enriched signal in basal zone Gαo neurons. Corresponding high-resolution images (right) show that the overall ER morphology at cellular reso-lution is indistinguishable between Cnpy1^+/+^ and Cnpy1^-/-^ at P14 age. Note that while cellular-scale organiza-tion is preserved, ultrastructural features, such as gyroid structures of the ER membrane are below the reso-lution limit for light microscopy.

**Figure S11.**
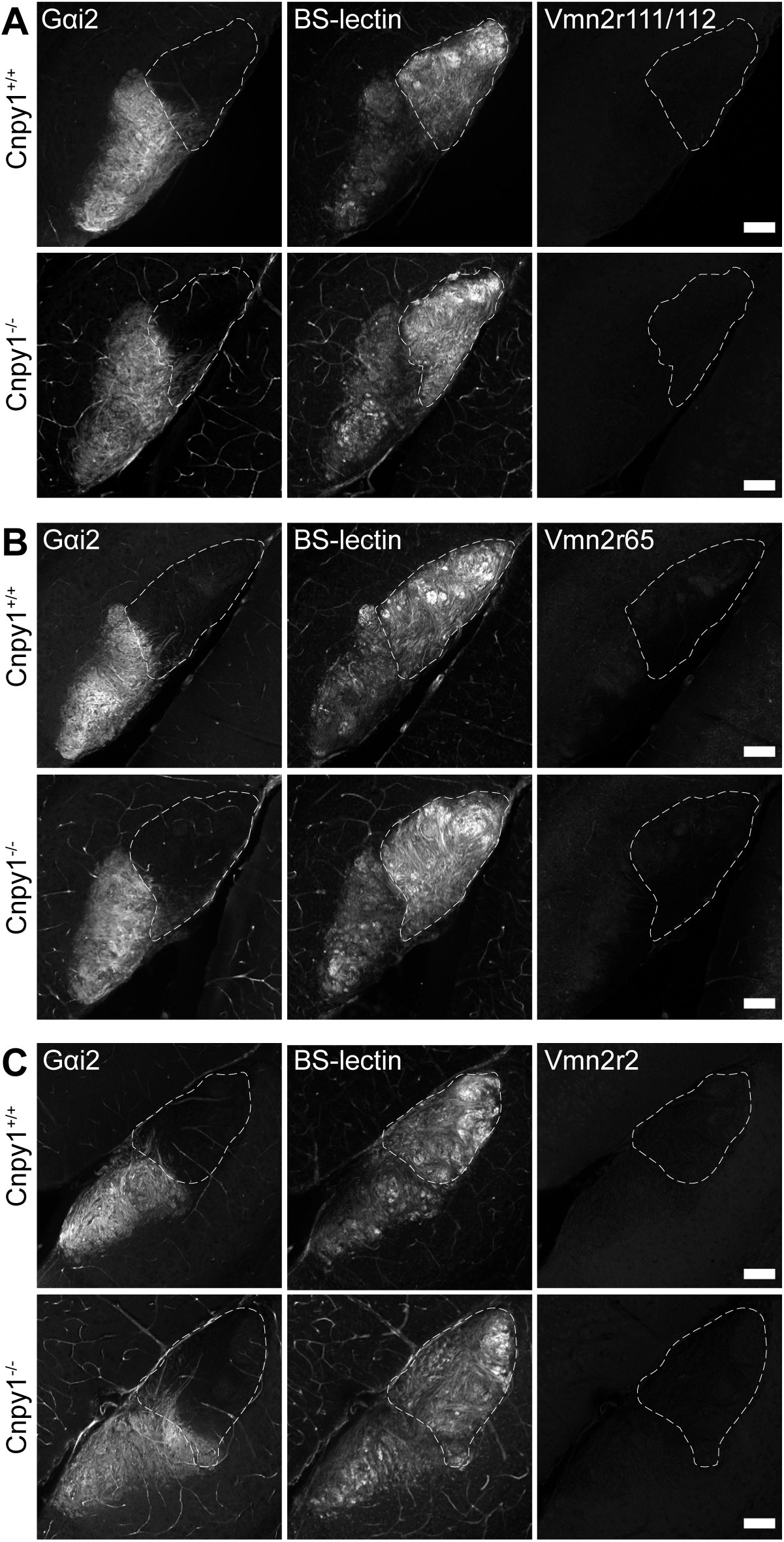
Absence of V2R localization to the AOB in Cnpy1^+/+^ as well as Cnpy1^-/-^ mice. **A-C)** Immuno-fluorescence labelling of anterior AOB marked with Gαi2 and posterior AOB preferentially labelled with BS-lectin along with V2Rs - Vmn2r111/112 (**A**), Vmn2r65 (**B**) and Vmn2r2 (**C**). Anti-V2R signal for these antibodies in the VNO is shown in Figure 3 and Figure S8; but was not observed in the AOB of either Cnpy1^+/+^ or Cnpy1^-/-^ at P14 age. Dashed line indicates the boundary of Gαo zone. Scale bar: 100 μm.

**Figure S12.**
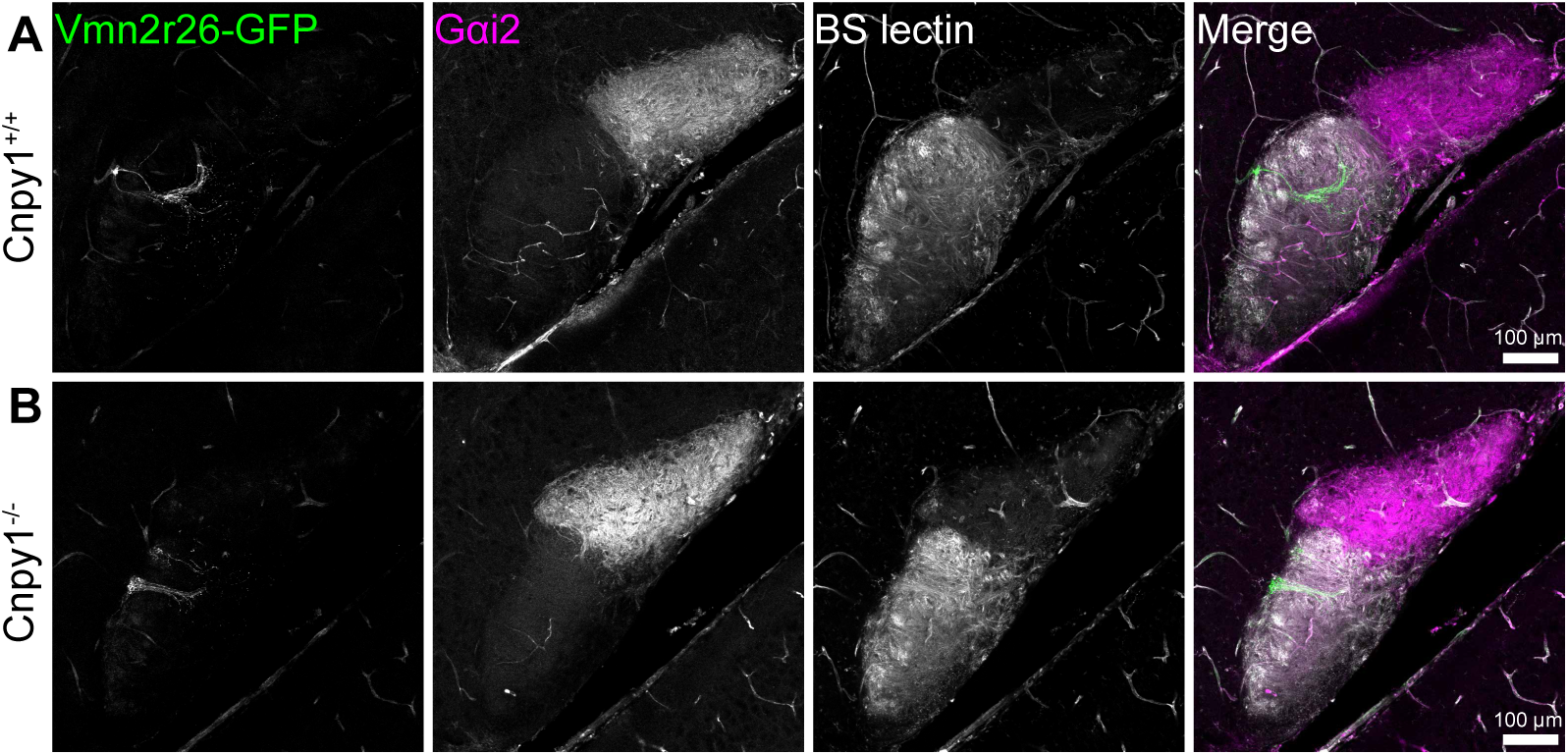
Normal targeting and glomerular coalescence of Vmn2r26-GFP axons in Cnpy1^-/-^ mice. **A-B**) Representative immunofluorescence confocal microscopy images of accessory olfactory bulb (AOB) sagittal sections obtained from P7 Vmn2r26-IRES-GFP, Cnpy1^+/+^ (**A**) or Vmn2r26-IRES-GFP, Cnpy1^-/-^ (**B**) AOB. Targeting and coalescence of GFP labeled projections to glomeruli within the posterior part of the AOB appear to be intact in the absence of Cnpy1. The anterior AOB is marked by Gαi2 and differentiated from posterior AOB which is preferentially labelled by BS-Lectin.

**Figure S13:**
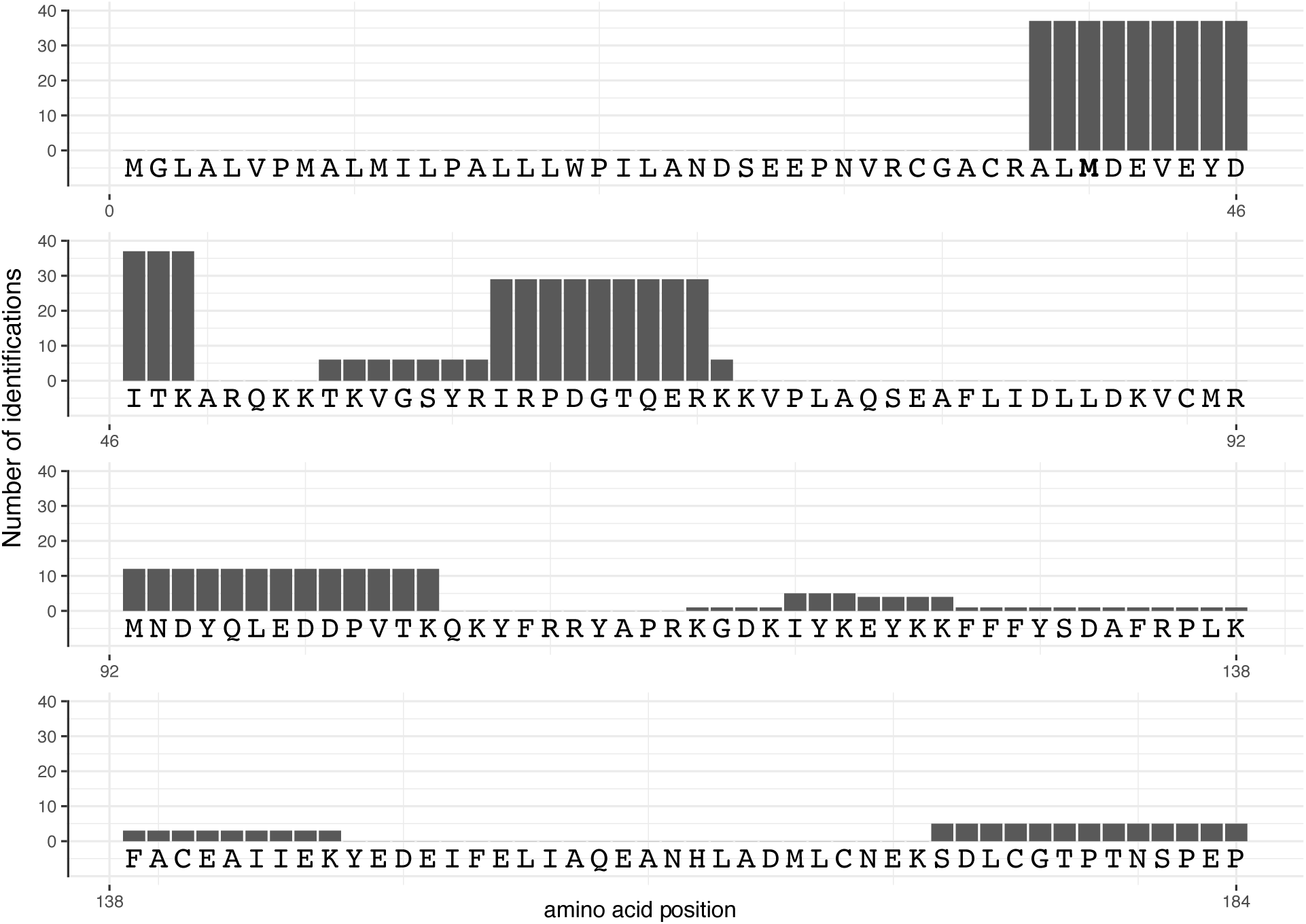
Alignment of Cnpy1 amino acid sequence with trypsin digested peptides detected by mass spec-trometry. Number of identifications for each peptide detected is marked on y-axis. Methionine (M40) in bold is the translation start site for Ensembl transcript - ENMUST00000141601. The presence of signal from a peptide containing AL before M40 re-confirms the upstream translation start site.

**Figure S14.**
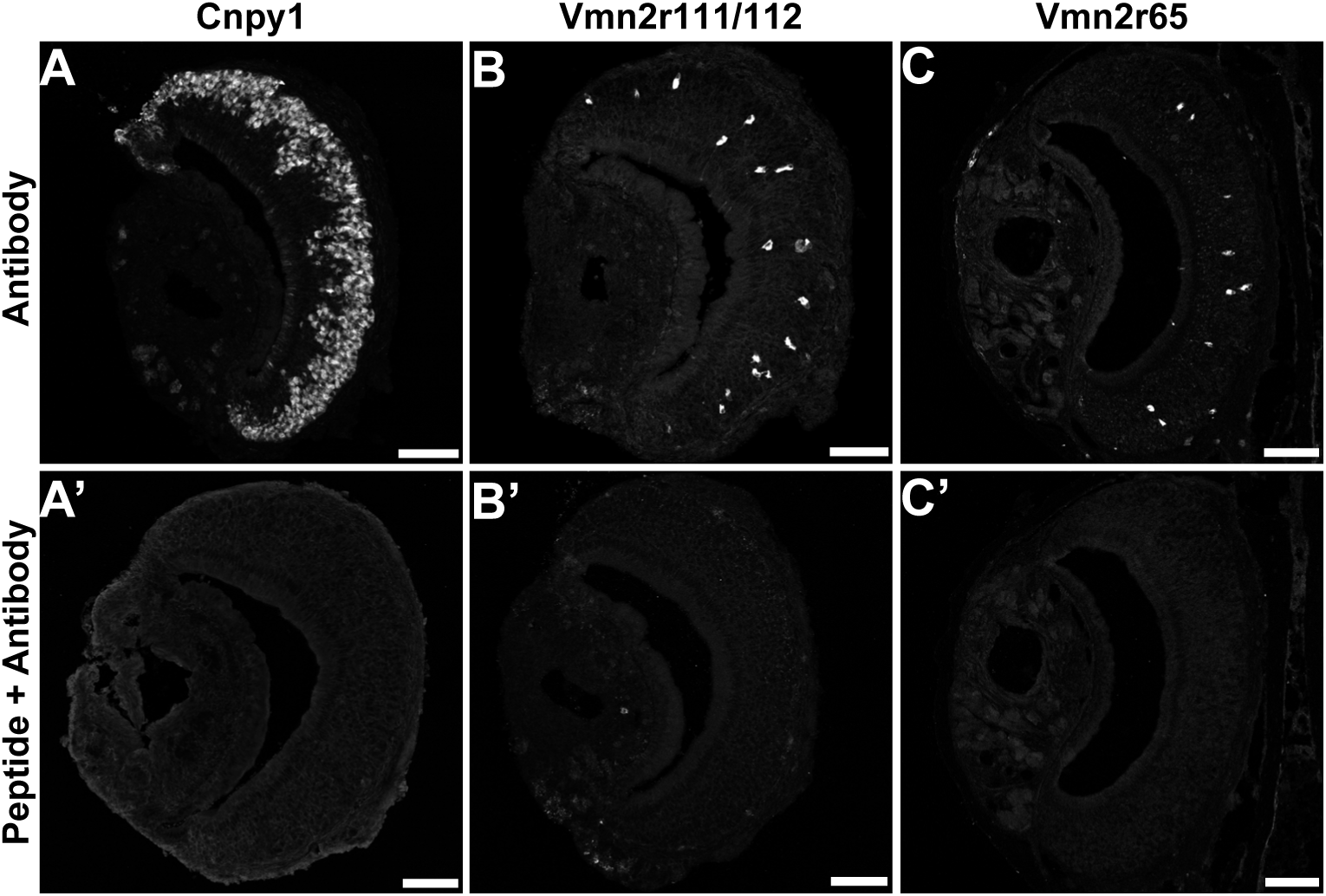
Generation of custom polyclonal antibodies. Anti-Cnpy1, anti-Vmn2r111/112 and anti-Vmn2r65 antibodies were generated by injecting KLH conjugated DDPVTKQKYFRRYAPRKGD, FWKMKRNENKDRNQ, and QKVNFTQKFSDTHSKIEYNH, respectively. Specific immunolabeling (**A-C**) with each antibody is blocked after pre-incubation of each antibody with the respective immunogenic peptide (**A’-C’**). Scale bar: 100 μm

**Figure S15.**
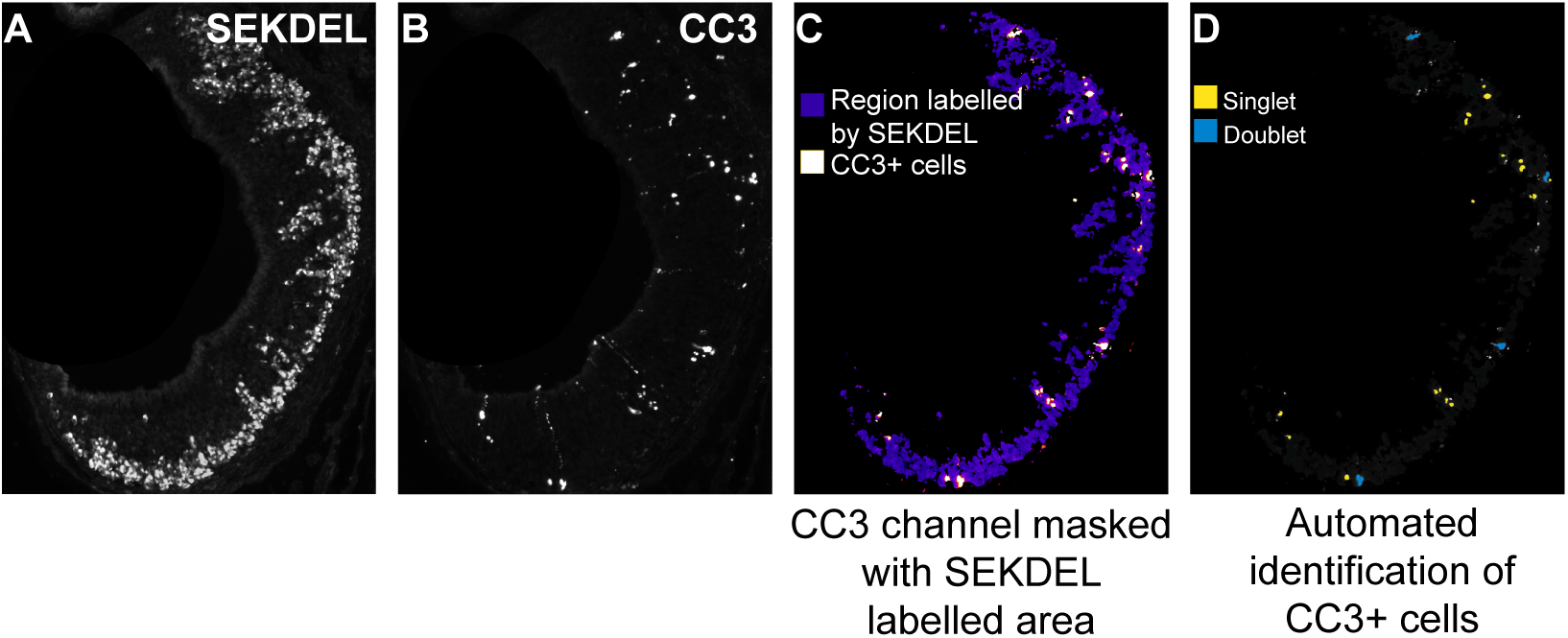
Automated quantification of cell numbers from images. **A-C**) Immunofluorescence images were manually masked to remove the non-sensory epithelium from all channels (**A, B**). An ilastik segmentation model was trained to identify pixels labeled by anti-SEKDEL (**A**) and to generate a mask, which was then applied to the CC3 channel (**B**) to segment the SEKDEL-marked region in CC-3 channel (**C**). **D**) An independently trained object identification model was subsequently used to detect CC3-positive cells, classifying them as either singlets or doublets within the SEKDEL-segmented CC3 images.

## References

1. S. D. Liberles, Mammalian pheromones. Annu Rev Physiol 76, 151–175 (2014).

2. P. Chamero et al., Identification of protein pheromones that promote aggressive behaviour. Nature 450, 899–902 (2007).

3. Y. Isogai et al., Molecular organization of vomeronasal chemoreception. Nature 478, 241–245 (2011).

4. H. Kimoto, S. Haga, K. Sato, K. Touhara, Sex-specific peptides from exocrine glands stimulate mouse vomeronasal sensory neurons. Nature 437, 898–901 (2005).

5. D. Lee, M. Kume, T. E. Holy, Sensory coding mechanisms revealed by optical tagging of physiologically defined neuronal types. Science 366, 1384–1389 (2019).

6. T. Ishii, J. Hirota, P. Mombaerts, Combinatorial coexpression of neural and immune multigene families in mouse vomeronasal sensory neurons. Curr Biol 13, 394–400 (2003).

7. J. Loconto et al., Functional expression of murine V2R pheromone receptors involves selective association with the M10 and M1 families of MHC class Ib molecules. Cell 112, 607–618 (2003).

8. N. D. Dwyer, E. R. Troemel, P. Sengupta, C. I. Bargmann, Odorant receptor localization to olfactory cilia is mediated by ODR-4, a novel membrane-associated protein. Cell 93, 455–466 (1998).

9. E. K. Baker, N. J. Colley, C. S. Zuker, The cyclophilin homolog NinaA functions as a chaperone, forming a stable complex in vivo with its protein target rhodopsin. EMBO J 13, 4886–4895 (1994).

10. E. E. Rosenbaum et al., XPORT-dependent transport of TRP and rhodopsin. Neuron 72, 602–615 (2011).

11. E. E. Rosenbaum, R. C. Hardie, N. J. Colley, Calnexin is essential for rhodopsin maturation, Ca2+ regulation, and photoreceptor cell survival. Neuron 49, 229–241 (2006).

12. H. Saito, M. Kubota, R. W. Roberts, Q. Chi, H. Matsunami, RTP family members induce functional expression of mammalian odorant receptors. Cell 119, 679–691 (2004).

13. S. Dey, H. Matsunami, Calreticulin chaperones regulate functional expression of vomeronasal type 2 pheromone receptors. Proc Natl Acad Sci U S A 108, 16651–16656 (2011).

14. R. Sharma et al., Olfactory receptor accessory proteins play crucial roles in receptor function and gene choice. Elife 6 (2017).

15. G. V. S. Devakinandan, M. Terasaki, A. Dani, Single-cell transcriptomics of vomeronasal neuroepithelium reveals a differential endoplasmic reticulum environment amongst neuronal subtypes. Elife 13 (2024).

16. Y. Hirate, H. Okamoto, Canopy1, a novel regulator of FGF signaling around the midbrain-hindbrain boundary in zebrafish. Curr Biol 16, 421–427 (2006).

17. D. Schildknegt et al., Characterization of CNPY5 and its family members. Protein Sci 28, 1276–1289 (2019).

18. S. T. Sowa et al., High-resolution Crystal Structure of Human pERp1, A Saposin-like Protein Involved in IgA, IgM and Integrin Maturation in the Endoplasmic Reticulum. J Mol Biol 433, 166826 (2021).

19. S. W. Santoro, S. Jakob, Gene expression profiling of the olfactory tissues of sex-separated and sex-combined female and male mice. Sci Data 5, 180260 (2018).

20. F. Teufel et al., SignalP 6.0 predicts all five types of signal peptides using protein language models. Nat Biotechnol 40, 1023–1025 (2022).

21. F. Hong et al., CNPY2 is a key initiator of the PERK-CHOP pathway of the unfolded protein response. Nat Struct Mol Biol 24, 834–839 (2017).

22. K. Konno et al., A molecule that is associated with Toll-like receptor 4 and regulates its cell surface expression. Biochem Biophys Res Commun 339, 1076–1082 (2006).

23. T. Shibata et al., PRAT4A-dependent expression of cell surface TLR5 on neutrophils, classical monocytes and dendritic cells. Int Immunol 24, 613–623 (2012).

24. Y. Shimizu, L. Meunier, L. M. Hendershot, pERp1 is significantly up-regulated during plasma cell differentiation and contributes to the oxidative folding of immunoglobulin. Proc Natl Acad Sci U S A 106, 17013–17018 (2009).

25. E. van Anken et al., Efficient IgM assembly and secretion require the plasma cell induced endoplasmic reticulum protein pERp1. Proc Natl Acad Sci U S A 106, 17019–17024 (2009).

26. Y. Wakabayashi et al., A protein associated with toll-like receptor 4 (PRAT4A) regulates cell surface expression of TLR4. J Immunol 177, 1772–1779 (2006).

27. T. Ishii, P. Mombaerts, Coordinated coexpression of two vomeronasal receptor V2R genes per neuron in the mouse. Mol Cell Neurosci 46, 397–408 (2011).

28. L. Silvotti, A. Moiani, R. Gatti, R. Tirindelli, Combinatorial co-expression of pheromone receptors, V2Rs. J Neurochem 103, 1753–1763 (2007).

29. J. M. Young, B. J. Trask, V2R gene families degenerated in primates, dog and cow, but expanded in opossum. Trends Genet 23, 212–215 (2007).

30. L. Silvotti, E. Cavalca, R. Gatti, R. Percudani, R. Tirindelli, A recent class of chemosensory neurons developed in mouse and rat. PLoS One 6, e24462 (2011).

31. K. Del Punta, A. Puche, N. C. Adams, I. Rodriguez, P. Mombaerts, A divergent pattern of sensory axonal projections is rendered convergent by second-order neurons in the accessory olfactory bulb. Neuron 35, 1057–1066 (2002).

32. I. Salazar, P. Sanchez Quinteiro, Differential development of binding sites for four lectins in the vomeronasal system of juvenile mouse: from the sensory transduction site to the first relay stage. Brain Res 979, 15–26 (2003).

33. J. H. Brann, S. Firestein, Regeneration of new neurons is preserved in aged vomeronasal epithelia. J Neurosci 30, 15686–15694 (2010).

34. H. Nakano et al., Activating transcription factor 5 (ATF5) is essential for the maturation and survival of mouse basal vomeronasal sensory neurons. Cell Tissue Res 363, 621–633 (2016).

35. R. R. Katreddi et al., Notch signaling determines cell-fate specification of the two main types of vomeronasal neurons of rodents. Development 149 (2022).

36. Y. Ohta, K. Ichimura, Proliferation markers, proliferating cell nuclear antigen, Ki67, 5-bromo-2’-deoxyuridine, and cyclin D1 in mouse olfactory epithelium. Ann Otol Rhinol Laryngol 109, 1046–1048 (2000).

37. G. E. Karagoz, D. Acosta-Alvear, P. Walter, The Unfolded Protein Response: Detecting and Responding to Fluctuations in the Protein-Folding Capacity of the Endoplasmic Reticulum. Cold Spring Harb Perspect Biol 11 (2019).

38. P. Walter, D. Ron, The unfolded protein response: from stress pathway to homeostatic regulation. Science 334, 1081–1086 (2011).

39. L. Belluscio, G. Koentges, R. Axel, C. Dulac, A map of pheromone receptor activation in the mammalian brain. Cell 97, 209–220 (1999).

40. I. Rodriguez, P. Feinstein, P. Mombaerts, Variable patterns of axonal projections of sensory neurons in the mouse vomeronasal system. Cell 97, 199–208 (1999).

41. S. Martini, L. Silvotti, A. Shirazi, N. J. Ryba, R. Tirindelli, Co-expression of putative pheromone receptors in the sensory neurons of the vomeronasal organ. J Neurosci 21, 843–848 (2001).

42. L. Silvotti, R. M. Cavaliere, S. Belletti, R. Tirindelli, In-vivo activation of vomeronasal neurons shows adaptive responses to pheromonal stimuli. Sci Rep 8, 8490 (2018).

43. P. Chamero et al., G protein G(alpha)o is essential for vomeronasal function and aggressive behavior in mice. Proc Natl Acad Sci U S A 108, 12898–12903 (2011).

44. H. Paek, M. W. Antoine, F. Diaz, J. M. Hebert, Increased beta-catenin activity in the anterior neural plate induces ectopic mid-hindbrain characteristics. Dev Dyn 241, 242–246 (2012).

45. B. E. Hart, R. I. Tapping, Cell surface trafficking of TLR1 is differentially regulated by the chaperones PRAT4A and PRAT4B. J Biol Chem 287, 16550–16562 (2012).

46. F. Hong et al., Mapping the Interactome of a Major Mammalian Endoplasmic Reticulum Heat Shock Protein 90. PLoS One 12, e0169260 (2017).

47. K. Takahashi et al., A protein associated with Toll-like receptor (TLR) 4 (PRAT4A) is required for TLR-dependent immune responses. J Exp Med 204, 2963–2976 (2007).

48. M. Rosenbaum et al., MZB1 is a GRP94 cochaperone that enables proper immunoglobulin heavy chain biosynthesis upon ER stress. Genes Dev 28, 1165–1178 (2014).

49. E. Xiong et al., MZB1 promotes the secretion of J-chain-containing dimeric IgA and is critical for the suppression of gut inflammation. Proc Natl Acad Sci U S A 116, 13480–13489 (2019).

50. M. Tanaka, H. Treloar, R. G. Kalb, C. A. Greer, S. M. Strittmatter, G(o) protein-dependent survival of primary accessory olfactory neurons. Proc Natl Acad Sci U S A 96, 14106–14111 (1999).

51. E. M. Norlin, F. Gussing, A. Berghard, Vomeronasal phenotype and behavioral alterations in G alpha i2 mutant mice. Curr Biol 13, 1214–1219 (2003).

52. L. Stowers, T. E. Holy, M. Meister, C. Dulac, G. Koentges, Loss of sex discrimination and male-male aggression in mice deficient for TRP2. Science 295, 1493–1500 (2002).

53. M. Lo et al., CNPY4 inhibits the Hedgehog pathway by modulating membrane sterol lipids. Nat Commun 13, 2407 (2022).

54. M. D. Paul et al., CNPY4 is a Lipid-Binding Regulator of Sphingolipid Homeostasis. bioRxiv 10.1101/2025.10.13.682155, 2025.2010.2013.682155 (2025).

55. H. J. Shayya et al., ER stress transforms random olfactory receptor choice into axon targeting precision. Cell 185, 3896–3912 e3822 (2022).

56. R. P. Dalton et al., Olfactory and Vomeronasal Receptor Feedback Employ Divergent Mechanisms of PERK Activation. bioRxiv 10.1101/239830, 239830 (2018).

57. M. Scordino et al., CNPY2 protects against ER stress and is expressed by corticostriatal neurons together with CTIP2 in a mouse model of Huntington’s disease. Front Mol Neurosci 17, 1473058 (2024).

58. F. A. Ran et al., Genome engineering using the CRISPR-Cas9 system. Nat Protoc 8, 2281–2308 (2013).

59. R. Behringer, Manipulating the mouse embryo : a laboratory manual (Cold Spring Harbor Laboratory Press, Cold Spring Harbor, New York, ed. Fourth edition., 2014), pp. xxii, 814 pages.

60. S. Pease, T. L. Saunders, International Society for Transgenic Technologies., Advanced protocols for animal transgenesis : an ISTT manual, Springer protocols (Springer, Heidelberg ; New York, 2011), pp. xv, 669 p.

61. A. Dobin et al., STAR: ultrafast universal RNA-seq aligner. Bioinformatics 29, 15–21 (2013).

62. M. Pertea et al., StringTie enables improved reconstruction of a transcriptome from RNA-seq reads. Nat Biotechnol 33, 290–295 (2015).

63. R. Patro, G. Duggal, M. I. Love, R. A. Irizarry, C. Kingsford, Salmon provides fast and bias-aware quantification of transcript expression. Nat Methods 14, 417–419 (2017).

64. M. I. Love, W. Huber, S. Anders, Moderated estimation of fold change and dispersion for RNA-seq data with DESeq2. Genome Biol 15, 550 (2014).

65. S. Berg et al., ilastik: interactive machine learning for (bio)image analysis. Nat Methods 16, 1226–1232 (2019).

66. J. Schindelin et al., Fiji: an open-source platform for biological-image analysis. Nat Methods 9, 676–682 (2012).

67. C. S. Hughes et al., Ultrasensitive proteome analysis using paramagnetic bead technology. Mol Syst Biol 10, 757 (2014).

68. C. S. Hughes et al., Single-pot, solid-phase-enhanced sample preparation for proteomics experiments. Nat Protoc 14, 68–85 (2019).

69. T. Werner et al., Ion coalescence of neutron encoded TMT 10-plex reporter ions. Anal Chem 86, 3594–3601 (2014).

70. A. T. Kong, F. V. Leprevost, D. M. Avtonomov, D. Mellacheruvu, A. I. Nesvizhskii, MSFragger: ultrafast and comprehensive peptide identification in mass spectrometry-based proteomics. Nat Methods 14, 513–520 (2017).

71. M. E. Ritchie et al., limma powers differential expression analyses for RNA-sequencing and microarray studies. Nucleic Acids Res 43, e47 (2015).

72. W. Huber, A. von Heydebreck, H. Sultmann, A. Poustka, M. Vingron, Variance stabilization applied to microarray data calibration and to the quantification of differential expression. Bioinformatics 18 **Suppl 1**, S96–104 (2002).

73. S. B. Yoon et al., Real-time PCR quantification of spliced X-box binding protein 1 (XBP1) using a universal primer method. PLoS One 14, e0219978 (2019).

